# High-throughput CRISPR screens to dissect macrophage-*Shigella* interactions

**DOI:** 10.1101/2020.04.25.061671

**Authors:** Yong Lai, Liang Cui, Gregory H. Babunovic, Sarah M. Fortune, John G. Doench, Timothy K. Lu

## Abstract

Shigellosis causes most diarrheal deaths worldwide, particularly affecting children. *Shigella* invades and replicates in the epithelium of the large intestine, eliciting inflammation and tissue destruction. To understand how *Shigella* rewires macrophages prior to epithelium invasion, we performed genome-wide and focused secondary CRISPR knockout and CRISPR interference (CRISPRi) screens in *Shigella flexneri*-infected human monocytic THP-1 cells. Knockdown of the Toll-like receptor 1/2 signaling pathway significantly reduced pro-inflammatory cytokine and chemokine production, enhanced host cell survival, and controlled intracellular pathogen growth. Knockdown of the enzymatic component of the mitochondrial pyruvate dehydrogenase complex enhanced THP-1 cell survival. Small molecule inhibitors blocking key components of these pathways had similar effects; these were validated with human monocyte-derived macrophages, which closely mimic the *in vivo* physiological state of cells post-infection. High-throughput CRISPR screens can elucidate how *S. flexneri* triggers inflammation and redirects host pyruvate catabolism for energy acquisition before killing macrophages, pointing to new shigellosis therapies.

## INTRODUCTION

Annually, there are more than one million cases of shigellosis (Kotloff et al., 2018). In 2016, over two hundred thousand people were killed by shigellosis globally (Khalil et al., 2018). More than 65% of these deaths occurred in children under 5 years old and in adults older than 70 years (Khalil et al., 2018), indicating that the fully developed, healthy human immune system may be sufficient to prevent and control *Shigella* infections. We therefore focused on immune cells to investigate susceptibility to this potentially lethal bacterium at the cellular level.

Infection is initiated when *Shigella* crosses the intestinal epithelium through microfold cells (M cells) (Wassef et al., 1989). After transcytosis to the M cell pocket, *Shigella* targets the resident macrophages, inducing caspase-1-dependent pyroptotic cell death, a step essential to subsequent invasion and replication in the intestinal epithelium (Ashida et al., 2014; Ashida et al., 2011). Epithelial cells constitute the major habitat of *Shigella* (Ashida et al., 2015). Within this replicative niche, *Shigella flexneri* delivers various virulence proteins via a type III secretion system (T3SS), resulting in weakened host defenses (Ashida et al., 2015; Killackey et al., 2016). These virulence proteins reduce intracellular trafficking (Ferrari et al., 2019; Ramel et al., 2011), antagonize caspase-4-dependent pyroptosis (Kobayashi et al., 2013), prevent necrosis mediated by mitochondrial damage (Carneiro et al., 2009), and inhibit the early stage of apoptosis by p53 degradation (Bergounioux et al., 2012). As a consequence, epithelial cells survive infection and continue to harbor the bacteria (Bergounioux et al., 2012; Niebuhr et al., 2002; Pendaries et al., 2006).

Whereas the effects of *Shigella* infection on epithelial cells have been studied extensively, little attention has been paid to how *S. flexneri* interacts with macrophages. Yet understanding these interactions is crucial to redirecting the immune response to shield against this bacterial infection. What we know so far is that this intracellular pathogen rapidly induces macrophage pyroptosis by activating inflammasomes, specifically NLRP3 (NLR family pyrin domain-containing protein 3) and NLRC4 (NLR family CARD domain-containing protein 4) (Suzuki et al., 2014a; Suzuki et al., 2007; Willingham et al., 2007). To improve our understanding of how *S. flexneri* manipulates macrophages and induces rapid cell death and to identify host targets for potential therapy, we conducted CRISPR screens in human macrophage-like THP-1 cells following infection with *S. flexneri* and validated our results with primary human monocyte-derived macrophages (hMDM).

## RESULTS

### Host cell survival-based genome-wide primary screens

To examine whether *S. flexneri* invades and kills THP-1 cells, we first assessed the phagocytosis of *S. flexneri* M90T by detecting the red fluorescence (*uhpT:: dsRed*) induced by host cell-produced glucose 6-phosphate (Runyen-Janecky and Payne, 2002) and indicative of intracellular *S. flexneri*. The reporter detects the presence of bacteria in the cytosol following their entry into the host cells (though once the bacteria have been taken up by endosomes, they are no longer detected). We then measured THP-1 cell viability post-infection (**Figure S1**). We found that *S. flexneri*, at a multiplicity of infection (MOI) of 10:1, efficiently infected THP-1 cells and induced host cell death 3 hours after infection (**Figures S1A, S1D and S1E**). This result indicated that infection with *S. flexneri* could be utilized as selective pressure for subsequent host survival-based genetic screens. Independent biological triplicates of genome-wide CRISPR knockout and CRISPRi screen libraries were prepared in THP-1 cells expressing Cas9 and dCas9-Krab, respectively (Doench et al., 2016; Lai et al., 2020; Sanson et al., 2018) (**Figure 1A**). After *S. flexneri* infection, surviving THP-1 cells with specific single guide RNA (sgRNA) barcodes were maintained in culture medium for continuous replication and harvested for next generation sequencing and analysis. The distribution of sgRNAs in *S. flexneri-infected* THP-1 cells was significantly different from that in uninfected THP-1 cells (**Figures S2A-S2F**). The results of genome-wide screens were visualized with volcano plots (**Figures 1B, 1C** and **S2G-S2L**). Pathway analysis identified both depleted and enriched biological processes in *S. flexneri-infected* THP-1 cells (**Figures 1D** and **1E**). Most of these biological pathways, many of which are essential for host cell functions, were depleted post-infection. Yet, CRISPR-Cas9 knockout and CRISPRi screens also identified several enriched biological pathways, such as Toll-like receptor (TLR) signaling and pyruvate metabolism pathways.

**Figure 1.**
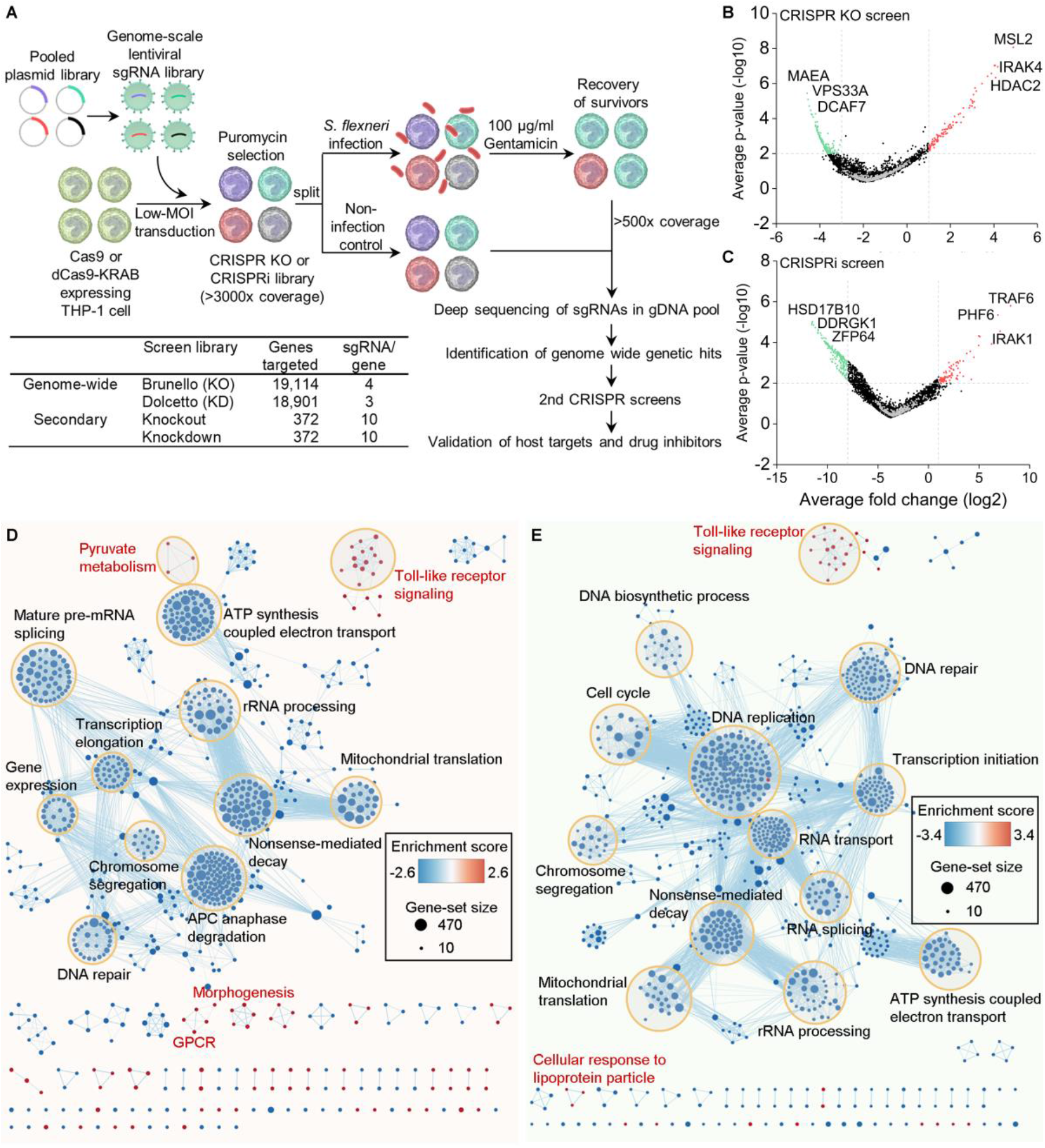
Genome-wide pooled CRISPR knockout and CRISPRi screens to dissect biological pathways in *S. flexneri* infection. (A) Monoclonal Cas9 and dCas9-Krab-expressing THP-1 cell lines were constructed and transduced with lentiviral sgRNA libraries. High coverage CRISPR knockout and CRISPRi libraries were split for subsequent *S. flexneri* infection. Surviving THP-1 cells with sgRNA barcodes were harvested and processed for next generation sequencing. Genome-wide genetic hits were identified by comparing sgRNA abundance between infected samples and non-infected controls. Based on those host targets, secondary CRISPR knockout and CRISPRi screen libraries were designed and prepared. Similarly, secondary positive screens were performed to validate those host targets. Finally, drug inhibitors that selectively inhibit genetic hits were tested. (B) and (C) Volcano plots from genome-wide CRISPR knockout (B) and CRISPRi (C) screens. For each sgRNA-targeted gene, the x axis shows its enrichment (positive hits) or depletion (negative hits) post-infection, and the y axis shows statistical significance measured by p-value. Top 3 positive and negative screen hits are labeled as red and green dots, respectively; positive hits being those that extended the survival of the THP-1 cells beyond 2-3 hours of bacterial infection and negative hits, those that shortened THP-1 cell survival. Gray dots represent non-targeting controls. For each screen, experiments were carried out in triplicate. (D) and (E), Genes identified by genome-wide CRISPR knockout (D) and CRISPRi (E) screens were functionally categorized to understand the biological functions involved in *S. flexneri* infection. Color gradient of nodes represents the enrichment scores of gene-sets. Node size represents the number of genes in the gene set. See also Figures S1 and S2.

In order to identify top positively selected genetic hits in *S. flexneri*-infected THP-1 cells, we used a false discovery rate (FDR) of <0.25 and log2 fold change of >1 as cut-off points. Positive hits were considered those that extended the survival of the THP-1 cells beyond 2-3 hours of bacterial infection. The CRISPR knockout screen, done in triplicate, identified more enriched genetic hits than the CRISPRi screens (**Figure 2A**). We observed positive selection of 73 and 28 genes in CRISPR-Cas9 knockout and CRISPRi screens, respectively, with 10 genes enriched in both screens (p-value of overlap <7.394E-18; **Figure 2A**). Pathway analysis identified multiple enriched biological processes in *S. flexneri-infected* THP-1 cells. Both CRISPR-Cas9 knockout and CRISPRi screens identified the same pathways, such as TLR cascades, pathways involved in chromatin organization and pyruvate metabolism, the cellular stress response pathway, and receptor tyrosine kinase signaling (**Figures S3A** and **S3B**; **Tables S1** and **S2**). Specifically, all key components of the TLR 1/2 signaling pathway were identified as positive hits in our genome-wide screens. These were: TRAF6 (TNF receptor-associated factor 6), IRAK1 (interleukin 1 receptor-associated kinase 1), IRAK4 (interleukin 1 receptor-associated kinase 4), MYD88 (myeloid differentiation primary response 88), TLR1, TLR2, TIFA (TRAF-interacting protein with forkhead-associated domain), and TIRAP (TIR domain-containing adaptor protein) (**Figures 2B** and **2C**; **Tables S3** and **S4**). Intriguingly, TRAF6 and TIFA also play important roles in the ALPK1 (alpha kinase 1)-TIFA-TRAF6-NF-κB pathway, by which epithelial cells detect lipopolysaccharide (LPS) biosynthetic intermediates and regulate inflammation in response to them (Garcia-Weber et al., 2018; Gaudet et al., 2017; Milivojevic et al., 2017). Yet, ALPK1, a newly identified cytosolic immune receptor in *Yersinia pseudotuberculosis* and *S. flexneri* infection (Garcia-Weber et al., 2018; Zhou et al., 2018), was not a genetic hit in our genome-wide CRISPR screens (**Figure 2B**). Moreover, several components of the type I interferon and the tumor necrosis factor (TNF) receptor signaling pathways, which induce pro-inflammatory cytokine and chemokine production or activate apoptotic cell death, were identified as positive hits; these included IFNAR1 (interferon alpha and beta receptor subunit 2), IFNAR2, TYK2 (tyrosine kinase 2), JAK1 (Janus kinase 1), STAT1 (signal transducer and activator of transcription 1), STAT2, IRF9 (interferon regulatory factor 9), TNFRSF1A (tumor necrosis factor receptor superfamily member 1A), and TNFRSF1B (**Figure 2C**). Although *S. flexneri* causes NF-κB-induced inflammation in both macrophages and epithelial cells, it also exhibits distinct mechanisms of host manipulation that may contribute to opposite outcomes of infection in these cell types, i.e., rapidly induced cell death of macrophages but inhibited cell death of epithelial cells (Ashida et al., 2015).

**Figure 2.**
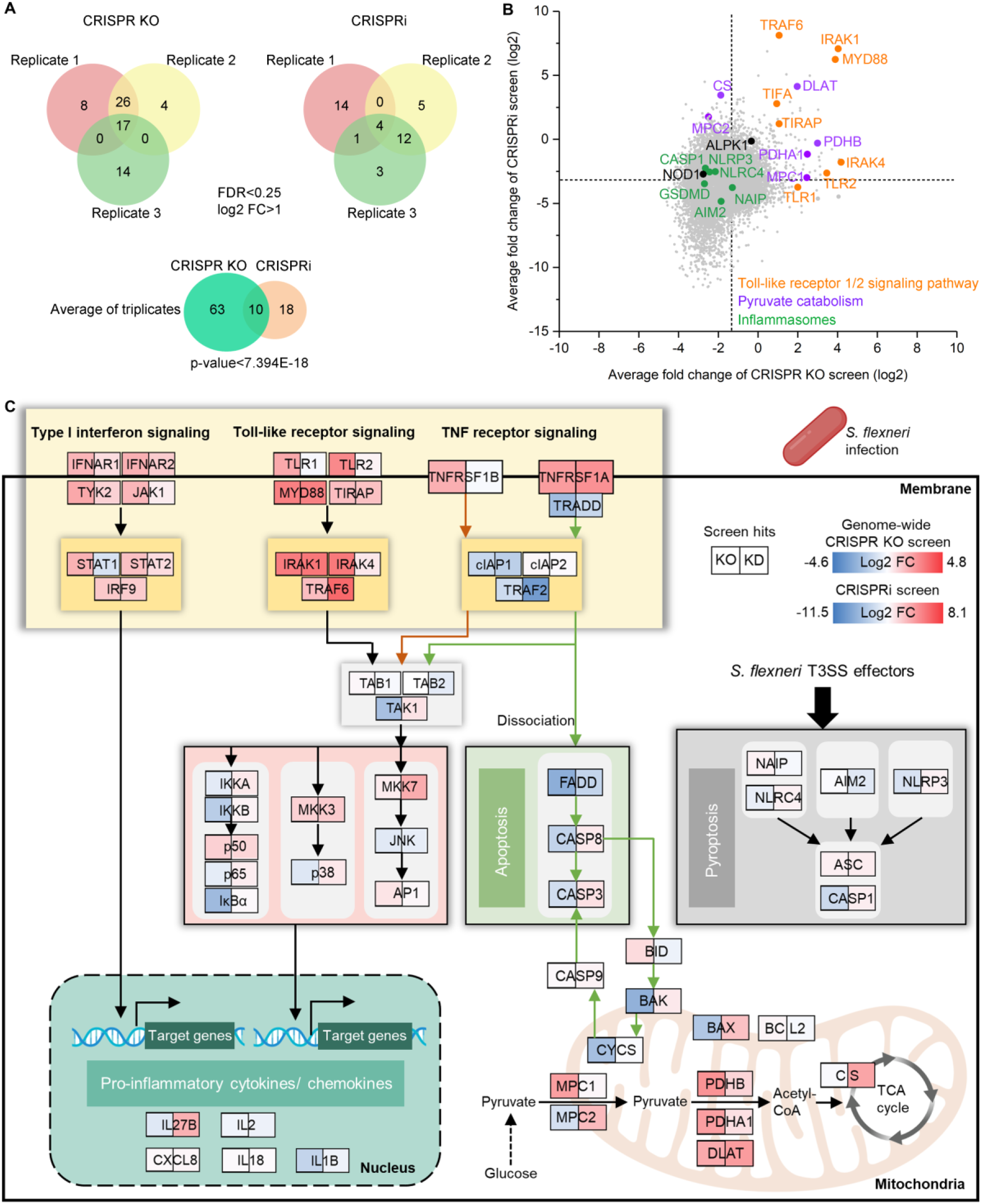
Genome-wide CRISPR knockout and CRISPRi screens to dissect enriched genes and biological pathways in *S. flexneri* infection. (A) Enriched genes in Venn diagram were filtered with a cut-off of FDR <0.25 and log2 fold change >1 in *S. flexneri* infection. The degree of significance of the overlap between genome-wide CRISPR knockout and CRISPRi screens is given. (B) Gene-centric visualization of average log2 fold change of CRISPR knockout and CRISPRi screens in *S. flexneri-infected* versus non-infected host cells. Selected components of TLR1/2, pyruvate catabolism signaling pathway, and inflammasome formation are highlighted in orange, purple, and green. (C) Top enriched genes and associated biological pathways in *S. flexneri* infection. Color gradient of gene boxes represents the log2 fold change of gene-sets in genome-wide CRISPR knockout and CRISPRi screens. See also Figure S3 and Tables S1-S4 and S7.

Moreover, the identification of genes as positive hits involved in pyruvate catabolism in macrophages, such as PDHB (pyruvate dehydrogenase E1 subunit beta), DLAT (dihydrolipoamide S-acetyltransferase), CS (citrate synthase), PDHA1 (pyruvate dehydrogenase E1 subunit alpha 1), MPC1 (mitochondrial pyruvate carrier 1), and MPC2 (**Figures 2B** and **2C**), echoed the rerouting of carbon flux by *S. flexneri* observed in epithelial (HeLa) cells (Kentner et al., 2014), which supports the rapid growth of intracellular bacteria using pyruvate as carbon source in the host cell (Waligora et al., 2014). Intriguingly, key components of the NLRC4 and NLRP3 inflammasomes, i.e., AIM2 (absent in melanoma 2), CASP1 (caspase 1), NLRC4, NLRP3, GSDMD (gasdermin D), and NAIP (NLR family apoptosis inhibitory protein), which are activated by *S. flexneri* T3SS effectors MxiI (Suzuki et al., 2014a), IpaB (Senerovic et al., 2012), and IpaH7.8 (Suzuki et al., 2014b) in macrophages, were identified in our genome-wide CRISPR screens as negative hits (i.e., a gene whose knockout or knockdown shortened THP-1 cell survival) (**Figures 2B** and **2C**), indicating that certain genes in the pyroptotic cell death pathway may actually protect host cells in *S. flexneri* infection. Overall, these results indicate the reliability of genome-wide CRISPR screens for studying *S. flexneri* infection and the importance of comprehensively understanding macrophage-*S. flexneri* interactions.

### Focused secondary screens for host-pathogen interactions

We next designed and prepared secondary CRISPR knockout and CRISPRi screen libraries targeting 372 human genes (Lai et al., 2020), in order to validate genome-wide screen hits, to test genes that were associated with different types of host cell death, and to compare the performance between CRISPR knockout and knockdown libraries by ensuring that there were consistent numbers of sgRNAs per gene in each type of library (**Figure 1A**). Similar to our genome-wide screens, surviving THP-1 cells were harvested following *S. flexneri* infection, and the results of the screens were visualized with volcano plots (**Figures S4A** and **S4B**). We identified 23 and 29 genes in secondary CRISPR knockout and CRISPRi screens, respectively, with 12 genes enriched in both screens (FDR<0.05; log2 FC>0.5; p-value<3.486E-09; **Figure 3A**). To evaluate the reliability of our genome-wide CRISPR screens, we calculated the validation rate of genes in secondary screens based on the FDR threshold (<5%); these genes were clustered by their p-value in the genome-wide screens (Sanson et al., 2018) (**Figure 3B**). The validation rate of screen hits in secondary CRISPR knockout and CRISPRi screens decreased with increasing p-values in primary genome-wide screens. This result suggests the reliability of genome-wide screens, which have fewer sgRNAs per gene than secondary screen libraries, for studying bacterial infections (**Figure 1A**). Furthermore, genome-wide top positive genetic hits were validated by secondary screens, such as genes in the TLR1/2 signaling pathway (IRAK1, MYD88, TRAF6), the type I interferon pathway (TYK2, IFNAR2, IRF8, STAT2), and the TNF receptor signaling pathway (TNFRSF1A), suggesting the robustness of genome-wide screens (**Figures 3C** and **S4C**; **Tables S5** and **S6**). Yet, genes involved in pyruvate catabolism, such as PDHB and PDHA1, were not scored significantly in secondary screens (**Figure S4C**); further validation of these genes would be required to confirm their involvement in the infectious process. Positive screen hits that were identified by both genome-wide and secondary screens were summarized based on the diverse signaling pathways they are associated with *S. flexneri* infection (**Figure 3D**). The advantage of using genome-wide screens to comprehensively identify host targets was indicated by the identification of many positive genetic hits with unknown functions in bacterial infection: PHF6 (PHD finger protein 6), PHIP (pleckstrin homology domain interacting protein), and TRERF1 (transcriptional regulating factor 1) (**Figure 3D** and **Tables S3** and **S4**).

**Figure 3.**
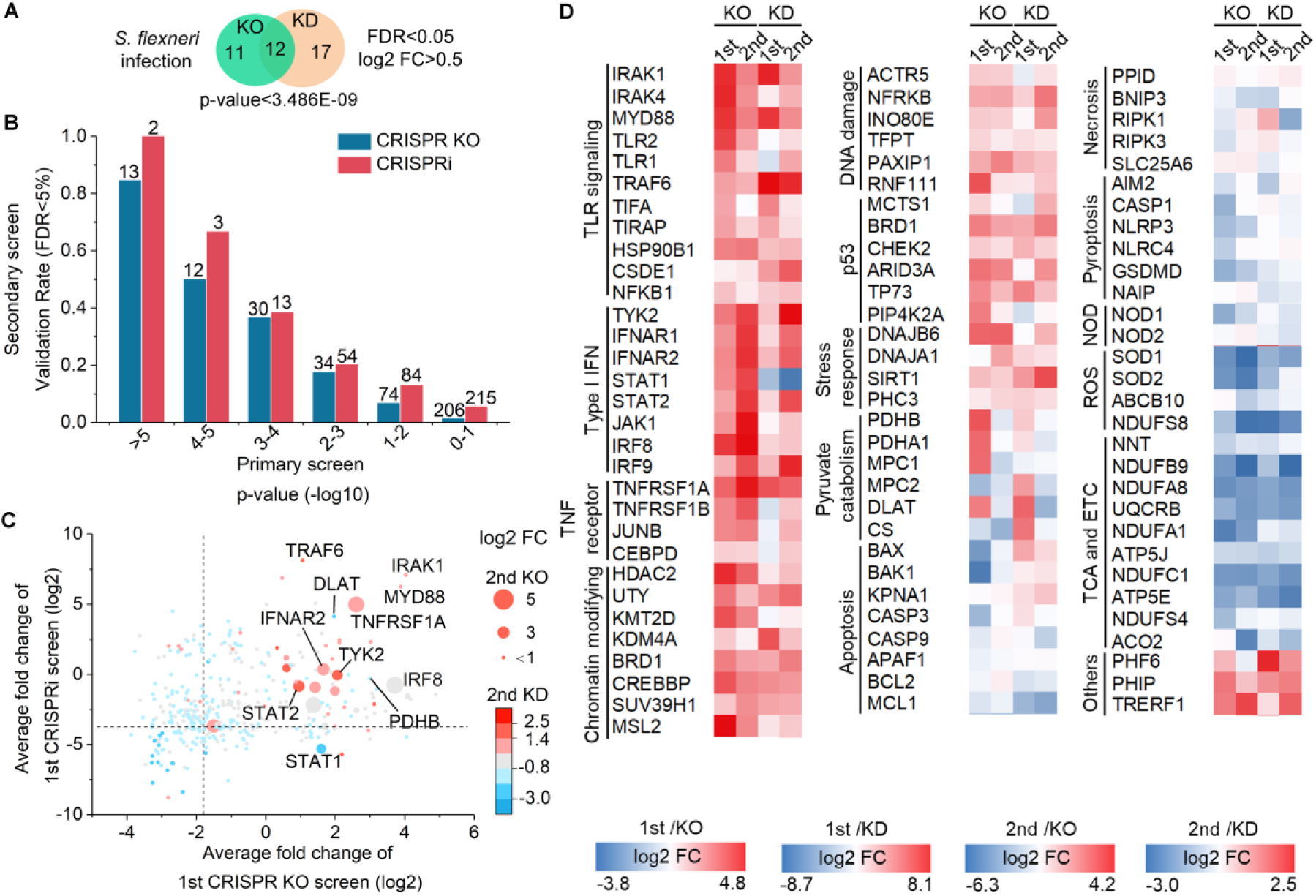
Secondary CRISPR knockout and CRISPRi screens identify host genetic hits in *S. flexneri* infection. (A) Enriched genes were filtered with a cut-off of FDR <0.05 and log2 fold change >0.5 in *S. flexneri* infection. The degree of significance of the overlap is given. (B) Validation rate of genetic hits in the secondary screen grouped by their p-value in the genome-wide screens in *S. flexneri* infection. Number of genes per category is indicated. (C) Genetic hits from both primary and secondary screens were ranked by their differential sgRNA abundance between *S. flexneri*-infected versus uninfected populations (log2 fold change). (D) Heatmap of screen hits clustered in different biological pathways in *S. flexneri* infection. See also Figures S4 and S5 and Tables S5 and S6.

Consistent with genome-wide CRISPR screens, components of inflammasomes mediating pyroptosis in macrophages post-infection, such as NLRP3 and GSDMD, were identified in our secondary screens as negative hits (**Figures 3D** and **S4C**). Yet, genes involved in other types of host cell death, i.e., necrosis and apoptosis, did not show consistent patterns of screen phenotypes and were not identified as genetic hits. Not surprisingly, NOD1 (nucleotide binding oligomerization domain containing 1), a critical intracellular bacterial sensor, was identified as a negative hit in the CRISPR knockout screen, indicating its protective role for host cells (**Figures 3D** and **S4C**). Intriguingly, SOD1 (superoxide dismutase 1) and SOD2, which destroy free superoxide radicals in host cells, were also identified as negative screen hits in *S. flexneri* infections (**Figures 3D** and **S4C**); decreased expression of these genes may contribute to necrosis induced by reactive oxygen species (He et al., 2011). Additionally, by calculating the sgRNA correlation of replicates, signal/noise ratio, p-value, and FDR of CRISPR screens, we found that CRISPR-Cas9 knockout and CRISPRi yielded comparable results in secondary screens (**Figure S5**).

### Screen hits validation for host-pathogen interactions

To verify the function of top positive genetic hits, we next constructed THP-1 cells with individual gene knockdowns and confirmed their phenotypes in *S. flexneri* infection. The positive correlation between screen phenotype and validation phenotype confirmed that repression of positive screen hits indeed enhanced host cell survival with a 92.3% true positive rate (Pearson R=0.56; **Figure 4A**). Moreover, repression of the transcription of MYD88, TRAF6, and IRAK1, key components in the TLR1/2 signaling pathway, also inhibited intracellular *S. flexneri* growth (**Figure 4B**).

**Figure 4.**
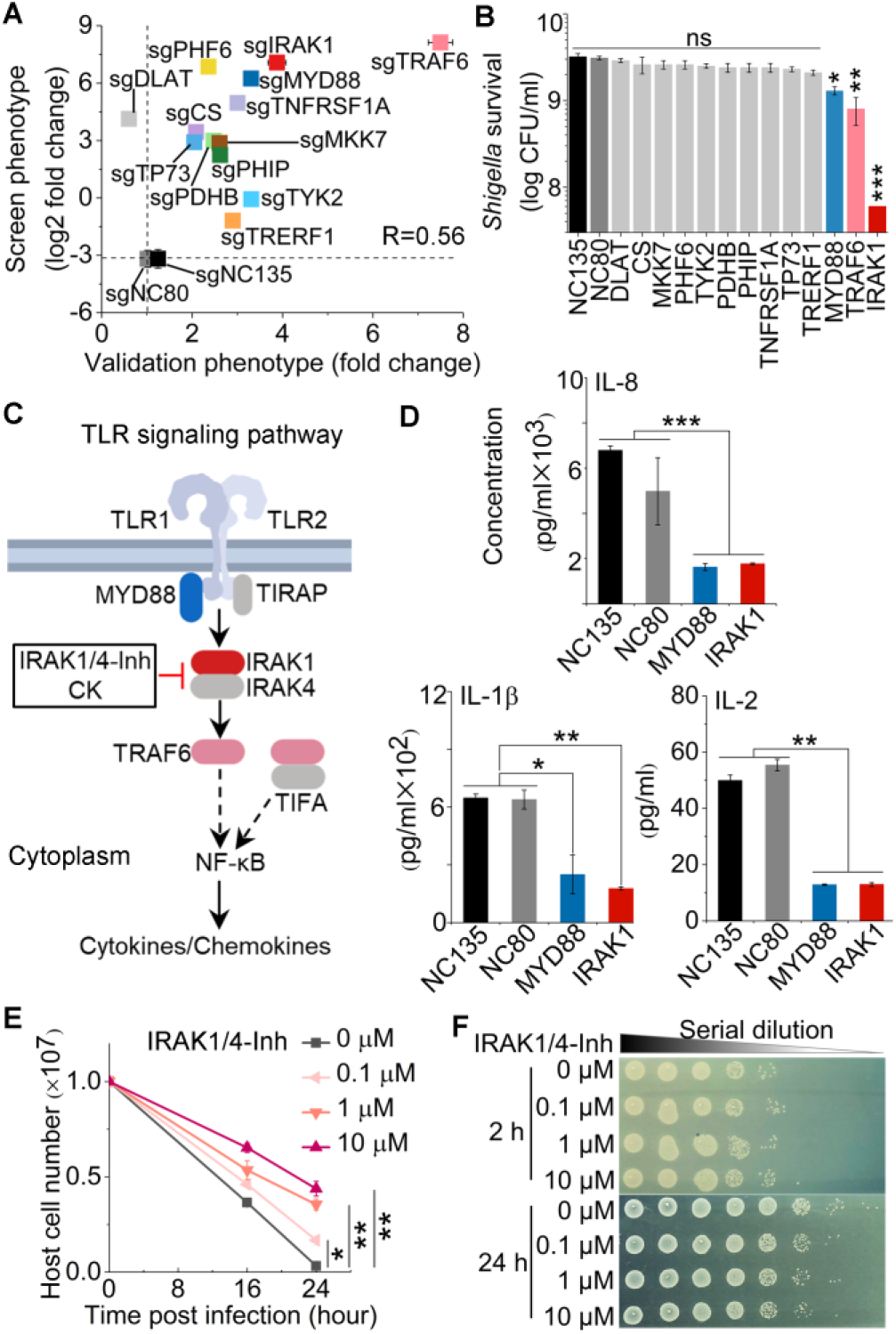
Validation of top positive genetic hits and effects of IRAK1 inhibitor in *S. flexneri* infection of human THP-1 cells. (A) Correlation between pooled screen and validation data. For each hit, the log2 fold change obtained from the genome-wide screening data (Screen phenotype) was plotted against the fold-change of cell viability of genetic hits compared to the non-targeting control cells (Validation phenotype). sgNC80 and sgNC135 are non-targeting controls. R is the Pearson correlation coefficient. (B) Intracellular *S. flexneri* level after infection of individual knockdown THP-1 cells. (C) Schematic of positive genetic hits in TLR1/2 signaling pathway and corresponding inhibitors. (D) Cytokine and chemokine production in infected THP-1 cells with MYD88 and IRAK1 knockdown. (E) and (F) The viability of THP-1 cells (E) and intracellular *S. flexneri* growth (F) post-infection in the presence or absence of IRAK1 inhibitor at different concentrations. Data represent the mean ±SD (n = 3) (two-tailed unpaired Student’s *t*-test, * P<0.05 ** P<0.01 *** P<0.001; ns represents not significant). See also Figures S6 and S7.

To characterize how the inhibition of those positive genetic hits mediates the host cell response and provides protection, we measured cytokine and chemokine production regulated by the TLR1/2 signaling pathway (**Figure 4C**). Knockdown of the transcription of either MYD88 or IRAK1 abolished the production of infection-induced pro-inflammatory cytokines and chemokines, such as IL-1β, IL-2, and IL-8 (**Figures 4D** and **S6A**). As a potential strategy to control intracellular bacterial infection by targeting host factors, we tested the function of corresponding small molecule inhibitors. IRAK1/4-Inh, a selective inhibitor of IRAK1, inhibited pro-inflammatory cytokine and chemokine production and protected host cells in a dose-dependent manner (**Figures 4E** and **S6B-S6D**). Compared to similar levels of intracellular *S. flexneri* after 2 hours of infection, the number of intracellular pathogens decreased in the presence of the IRAK1 inhibitor 24 hours post-infection, indicating that inhibition occurred by controlling intracellular pathogen growth rather than by blocking pathogen entry (**Figure 4F**). Ginsenoside Compound K (CK), a metabolite of Panax ginseng that also inhibits IRAK1 (Wee et al., 2015), similarly enhanced host cell survival, inhibited *S. flexneri* growth, and abolished infection-induced IL-8 production in THP-1 cells (**Figure S7**).

In addition to dysregulating the host immune response, *S. flexneri* grows rapidly and replicates in host cells but does so only if there is an adequate supply of nutrients. Knockout or knockdown of components of the pyruvate dehydrogenase complex or the pyruvate transporter MPC1/2 in the mitochondria redirected central metabolism, favoring the survival of THP-1 cells infected with *S. flexneri* (**Figures 3D** and **5A**), which is congruent with the induction by *S. flexneri* in epithelial cells of the production of acetyl-CoA (Kentner et al., 2014). We next tested the function of the PDHB inhibitor oxythiamine (OT), as well as its combination with an IRAK1 inhibitor (IRAK1/4-Inh), in *S. flexneri* infection. OT treatment enhanced host cell survival post-infection (**Figure 5B**) but failed to control intracellular *S. flexneri* growth (**Figure 5C**), which is consistent with the PDHB gene knockdown phenotype (**Figures 4A** and **4B**). Interestingly, the combination of both IRAK1 and PDHB inhibitors (IRAK1/4-Inh and OT) significantly enhanced host cell survival and controlled *S. flexneri* growth better than treatment with either of these inhibitors alone, indicating a synergistic effect of inhibitors targeting both immune and non-immune pathways in macrophages (**Figures 5B** and **5C**). Furthermore, in a PMA-stimulated THP-1-*S. flexneri* infection model, IRAK1/4-Inh, OT, and their combination enhanced host cell survival and limited intracellular pathogen growth (**Figures 5D-5F**).

**Figure 5.**
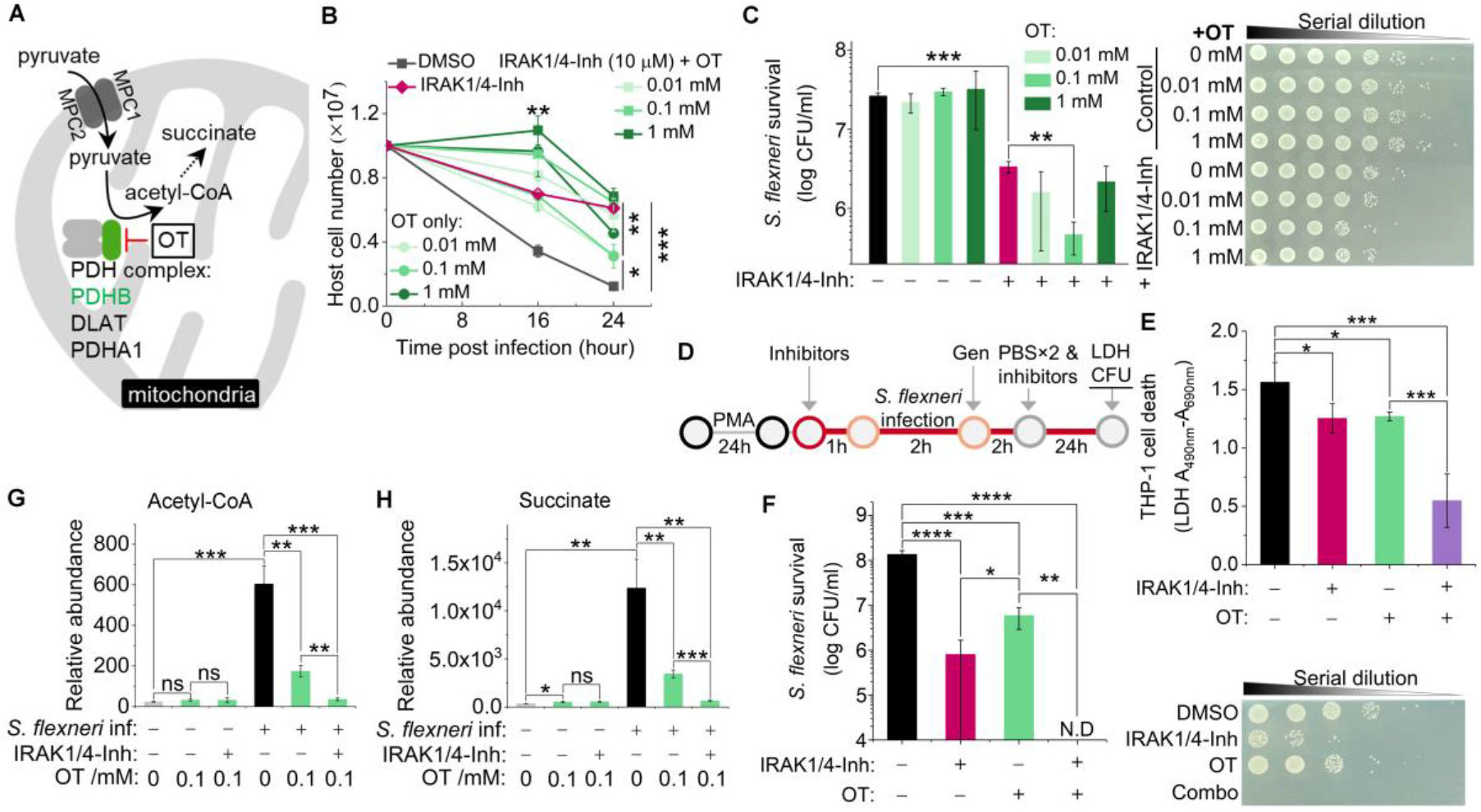
Validation of positive genetic hits in pyruvate catabolism signaling pathway and effects of corresponding inhibitor in *S. flexneri* infection of human THP-1 cells. (A) Schematic of positive genetic hits in pyruvate catabolism signaling pathway and corresponding inhibitor. (B) and (C) The growth of THP-1 cells (B) and intracellular *S. flexneri* level (C) post-infection in the presence or absence of various concentrations of PDHB inhibitor (oxythiamine, OT) or OT combined with IRAK1/4-Inh (10 μM). (D) Schematic of inhibitor validation in PMA-stimulated THP-1 cell infected with *S. flexneri*. (E) Effects of IRAK1 and PDHB inhibitors on the survival of differentiated THP-1 post-infection, with death measured by lactate dehydrogenase (LDH) release. (F) Effects of IRAK1 and PDHB inhibitors on the growth of intracellular *S. flexneri* in differentiated THP-1 cells. IRAK1/4-Inh is used at 10 μM. OT is used at 0.1 mM. (G) and (H) Production of acetyl-CoA (G) and succinate (H) with or without *S. flexneri* infection and in the presence or absence of OT or OT combined with IRAK1 inhibitor (10 μM). Data represent the mean ± SD (B, C, G and H, n = 3; E and F, n = 4) (two-tailed unpaired Student’s *t*-test, * P<0.05 ** P<0.01 *** P<0.001 **** P<0.0001; ns represents not significant; N.D represents not detectable).

In line with a previous study of epithelial cells (Kentner et al., 2014), we found that *S. flexneri* induced acetyl-CoA production in THP-1 cells, suggesting that, in both cases, *S. flexneri* supports its own rapid intracellular growth and replication by manipulating the central metabolism of the host cell (**Figure 5G**). Moreover, 0.1 mM of PDHB inhibitor decreased infection-induced acetyl-CoA and downstream succinate production, which shifts host metabolism and leads to enhanced host cell survival (**Figures 5G** and **5H**). The combination of both IRAK1 and PDHB inhibitors reduced acetyl-CoA and succinate production to the uninfected levels, thus limiting intracellular *S. flexneri* growth and propagation (**Figures 5G** and **5H**).

To further validate the function of inhibitors, we tested host cell death and intracellular *S. flexneri* growth in a primary human macrophage infection model (**Figure 6A**). Consistent with the results in the THP-1 infection model, IRAK1/4-Inh, OT, and their combination each significantly reduced the death of hMDMs by intracellular *S. flexneri* as measured by lactate dehydrogenase (LDH) release (**Figure 6B**). Those small molecule inhibitors also restricted intracellular *S. flexneri* growth in these cells as measured by counting bacterial colony forming units (CFUs) (**Figure 6C**). Thus, inhibiting the TLR 1/2 signaling pathway or the pyruvate catabolism signaling pathway restricts the intracellular pathogen burden and preserves the survival of human macrophages infected with *S. flexneri*.

**Figure 6.**
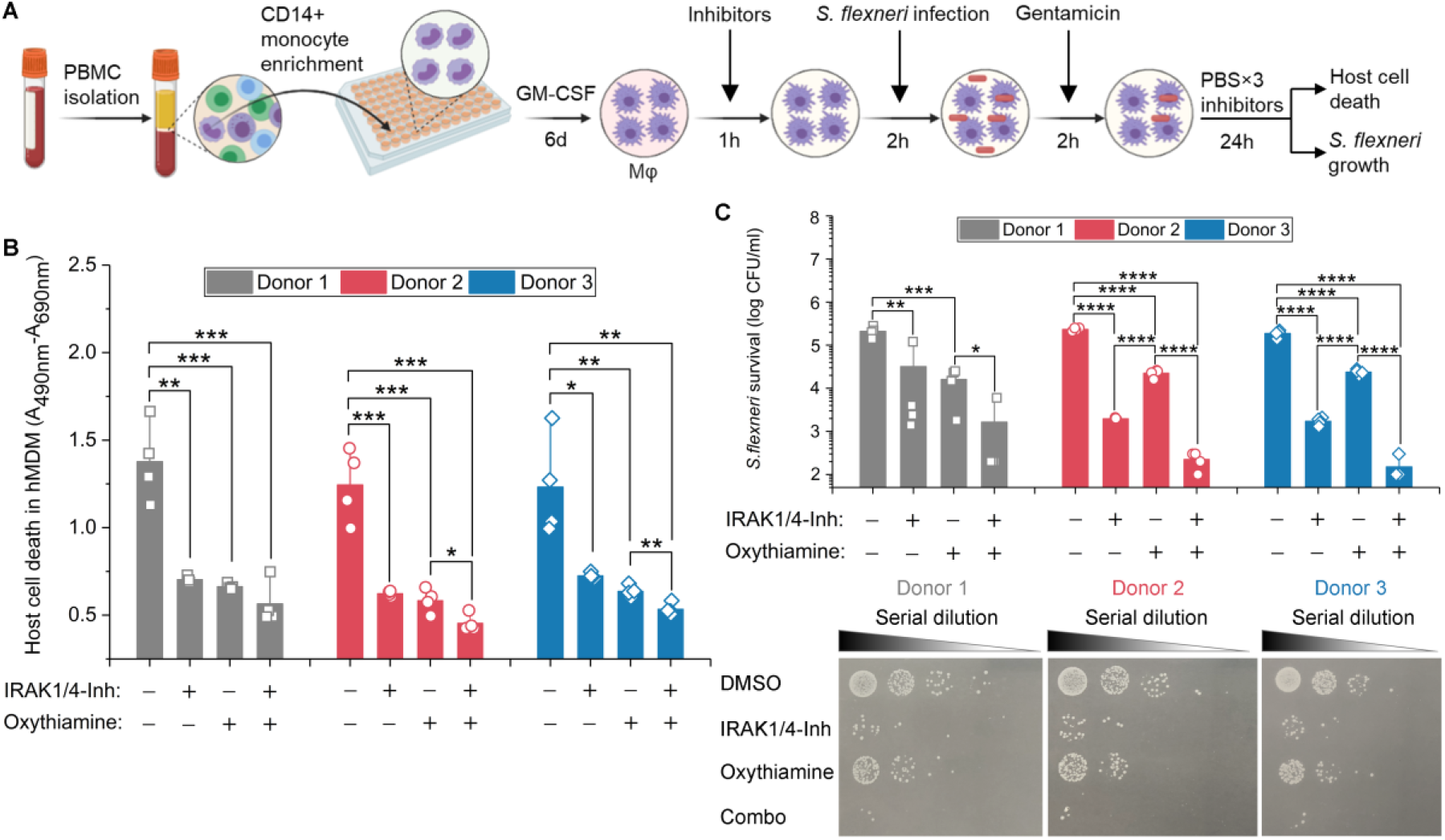
Validation of inhibitors in *S. flexneri* infection of primary human macrophages. (A) Schematic of inhibitor validation in *S. flexneri* infection cell models. (B) Effects of IRAK1 and PDHB inhibitors on the survival of primary human monocyte-derived macrophages (hMDM) infected with *S. flexneri*, with host cell death measured by LDH release. (C) Effects of IRAK1 and PDHB inhibitors on the growth of intracellular *S. flexneri* in hMDM, as measured by CFU. IRAK1/4-Inh was used at 10 μM. Oxythiamine (OT) was used at 0.1 mM. Data represent the mean ±SD (n = 4) (two-tailed unpaired Student’s t-test, * P<0.05 ** P<0.01 *** P<0.001 **** P<0.0001).

## DISCUSSION

Our study highlights the capability of host cell survival-based CRISPR screens to elucidate complex macrophage-pathogen interactions and to identify key cellular processes that are disrupted by intracellular pathogens. It thus reinforces the value of CRISPR screens to understand pathogenesis (Daniloski et al., 2020; Wei et al., 2020). Also highlighted is the importance of screening gene perturbations in the specific cell types that are infected by an intracellular pathogen, especially in the instance of intracellular bacteria, such as *Shigella*, that invade and inhabit more than one type of human cell.

Rapid macrophage death is prerequisite for *S. flexneri* to infect and persist in epithelial cells, which ultimately results in diarrhea and even dysentery, the most life-threatening manifestations of infection. However, unlike the intensive studies of the effects of *Shigella* infection on epithelial cells, the comprehensive interactions between *S. flexneri* and macrophages have been largely overlooked (Senerovic et al., 2012; Suzuki et al., 2014a; Suzuki et al., 2014b; Suzuki et al., 2007; Willingham et al., 2007). In epithelial cells, NF-κB-related inflammatory signaling is one of the major defenses against *S. flexneri* infection (Garcia-Weber et al., 2018; Milivojevic et al., 2017). For instance, upon sensing ADP-β-D-manno-heptose (ADP-Hep), epithelial cells activate NF-κB signaling in their cytosol and produce the pro-inflammatory chemokine IL-8 (Garcia-Weber et al., 2018). In response to these signals, *S. flexneri* produces multiple virulence proteins, disrupting inflammation and preventing epithelial cell death (Killackey et al., 2016). However, what occurs in epithelial cells does not necessarily occur in other cell types. In fact, our study revealed effects of *S. flexneri* in macrophage-like THP-1 cells and primary human macrophages distinct from those reported for *Shigella*-infected epithelial cells.

In THP-1 cells, *S. flexneri* also stimulated IL-8 production but did so by activating the TLR1/2 signaling pathway, and infection induced rapid THP-1 cell death. Although the TLR1/2 pathway is well known for its role in the innate immune response to invading pathogens via the recognition of peptidoglycan and triacyl lipopeptides, the induced inflammation could contribute to bacterial pathogenesis (Oliveira-Nascimento et al., 2012). For instance, in *Burkholderia* infection, knockout of TLR2 enhances the survival of mice and reduces sepsis, compared to what was observed with wild-type mice (Wiersinga et al., 2007). Moreover, the blockade of both TLR2 and TLR4 with monoclonal antibodies effectively inhibits the immunopathology triggered by *Escherichia coli* and *Salmonella enterica* and prevents mouse death (Spiller et al., 2008). In our study, inhibiting the TLR1/2 signaling pathway in THP-1 cells by IRAK1 inhibitors reduced pro-inflammatory cytokine and chemokine production, enhanced the survival of both THP-1 cells and primary human macrophages, and limited intracellular *S. flexneri* growth and replication, indicating the detrimental effect of the TLR1/2 signaling pathway on innate immune cells during *S. flexneri* infection. Besides, we found that host cell targets of *S. flexneri* virulence factors which had been identified in epithelial cells were not identified by our screens as positive genetic hits in THP-1 cells (**Table S7**). Considering the opposite effects of NF-κB signaling when epithelial cells and immune cells are infected with *S. flexneri*, modulating the inflammatory response of the host as a therapeutic strategy should be very carefully considered.

It is well known that macrophage pyroptosis is triggered by *S. flexneri*, allowing bacteria to escape from macrophages and invade epithelial cells (Ashida et al., 2015). However, this process also restricts intracellular bacterial pathogens by cytokine-independent mechanisms in a mouse model (Miao et al., 2010) or by the direct antibacterial effect of GSDMD-NT (the N-terminal cleavage product of GSDMD in pyroptosis) (Liu et al., 2016). It has been unclear whether pyroptosis protects the macrophages or the intracellular bacteria. Our host genetic perturbation strategies provide direct causal evidence that some of the genes that contribute to pyroptosis actually benefit the host cells, since knockout or knockdown of key components of pyroptosis (GSDMD and NLRP3) decreased host cell survival post-infection (**Figures 3D** and **S4C**).

Treatment for shigellosis is becoming increasingly difficult as resistance to most inexpensive and widely used antibiotics becomes more prevalent (Gu et al., 2012). In order to reduce mortality from diarrhea in children under 5 years of age to less than 1/1000 live births by 2025 (WHO and UNICEF, 2013), current antibiotics will have to be complemented by other kinds of treatment, such as host-directed therapies. Given that macrophages and epithelial cells appear to be manipulated by *S. flexneri* in diametrically opposite ways, developing an adjuvant therapy by targeting a common feature of those two types of host cells may be one way to block bacterial pathogenesis. In this study, we demonstrated that by inhibiting infection-induced acetyl-CoA production in host immune cells, the function of these cells can be restored and energy acquisition by the intracellular pathogen can be limited (**Figures 5** and **6**). Future studies will involve extensive investigation of more host targets and biological pathways, especially those with unknown functions in bacterial infection. In summary, our study not only sheds new light on the mechanisms underlying macrophage-*S. flexneri* interactions and *Shigella* pathogenesis but also provides new insights to guide the development of adjuvant therapy for shigellosis treatment.

## ACKNOWLEDGMENTS

We thank Karen Pepper for editing the manuscript. We thank Prof. Peter Dedon for reviewing the manuscript. *Shigella flexneri* M90T *ΔvirG* pCK100 (P*uhpT*::dsRed) was a gift from Cecile Arrieumerlou, Institut Cochin, France. This work was supported by the United States Defense Threat Reduction Agency (HDTRA1-15-1-0050 to T.K.L.), and the National Research Foundation, Prime Minister’s Office, Singapore under its Campus for Research Excellence and Technological Enterprise (CREATE) (Y.L., L.C., and T.K.L.).

## AUTHOR CONTRIBUTIONS

Y.L., L.C., G.H.B, S.M.F, J.G.D., and T.K.L. conceived and designed the research; Y.L. and J.G.D. performed and analyzed genome-wide CRISPR screen experiments; Y.L. and J.G.D. designed, performed and analyzed secondary CRISPR screen experiments; Y.L. and G.H.B conducted validation experiments of genetic hits and small molecule inhibitors; Y.L. and L.C. designed and performed metabolite profile experiments; Y.L. and T.K.L. coordinated the overall research; T.K.L. supervised the overall research; Y.L., L.C., G.H.B, S.M.F, J.G.D., and T.K.L. analyzed the data and wrote the manuscript. All authors discussed the results and reviewed the paper.

---

**Figure S1.**
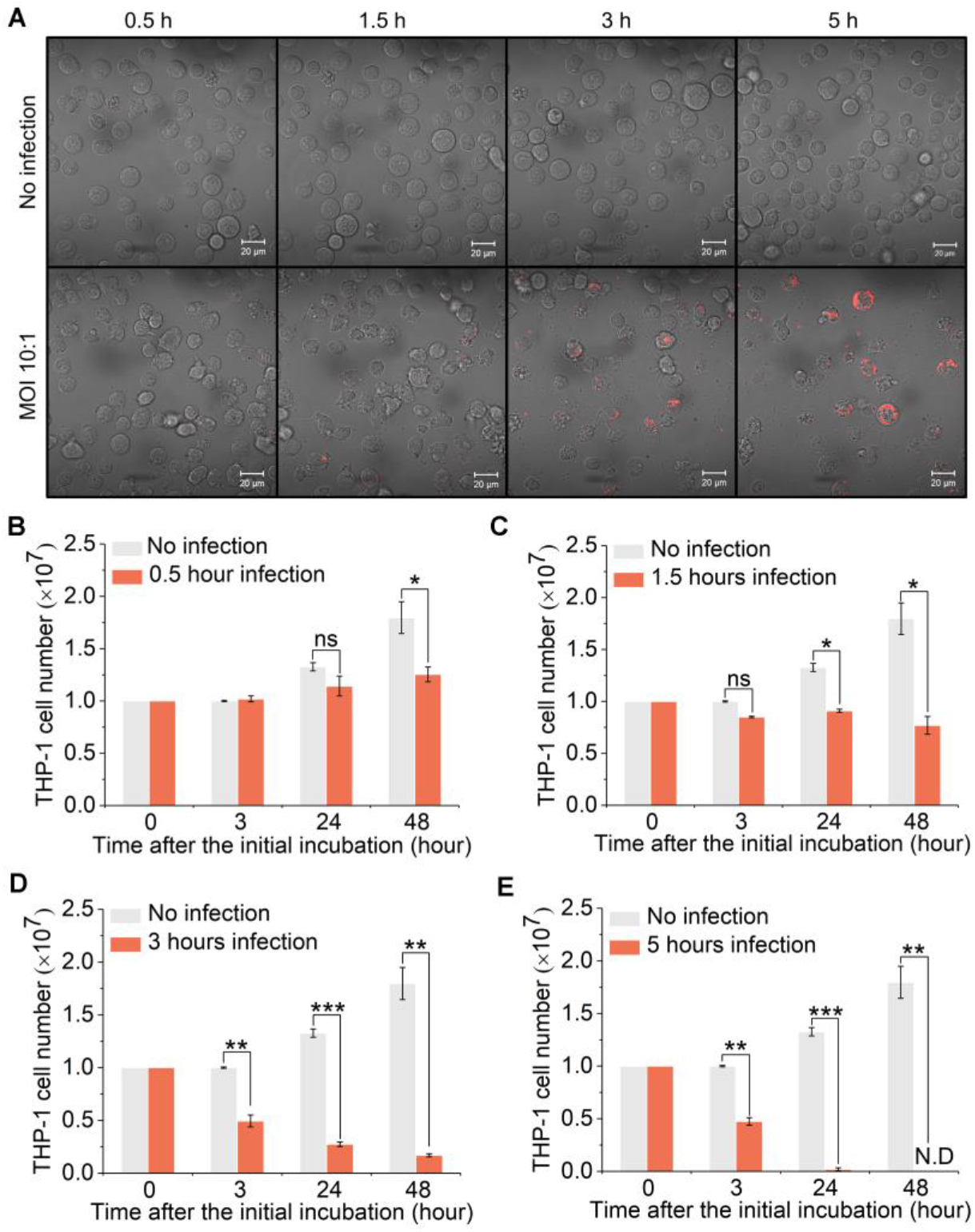
Optimization of conditions for *Shigella* infection of THP-1 cells. Related to Figure 1. (A) *S. flexneri* infection at an MOI (multiplicity of infection, or number of bacterial cells per host cell) of 10, from 0.5 hour to 5 hours incubation with THP-1 cells (Scale bar, 20 μm). (B)-(E) The number of *Shigella*-infected THP-1 cells after 0.5 to 5 hours infection. More than 90% of host cells were killed after 3 hours of infection (D). N.D represents not detectable. Data represent mean ±SD (n = 3) (two-tailed unpaired Student’s *t*-test, * P<0.05 ** P<0.01 *** P<0.001; ns represents not significant).

**Figure S2.**
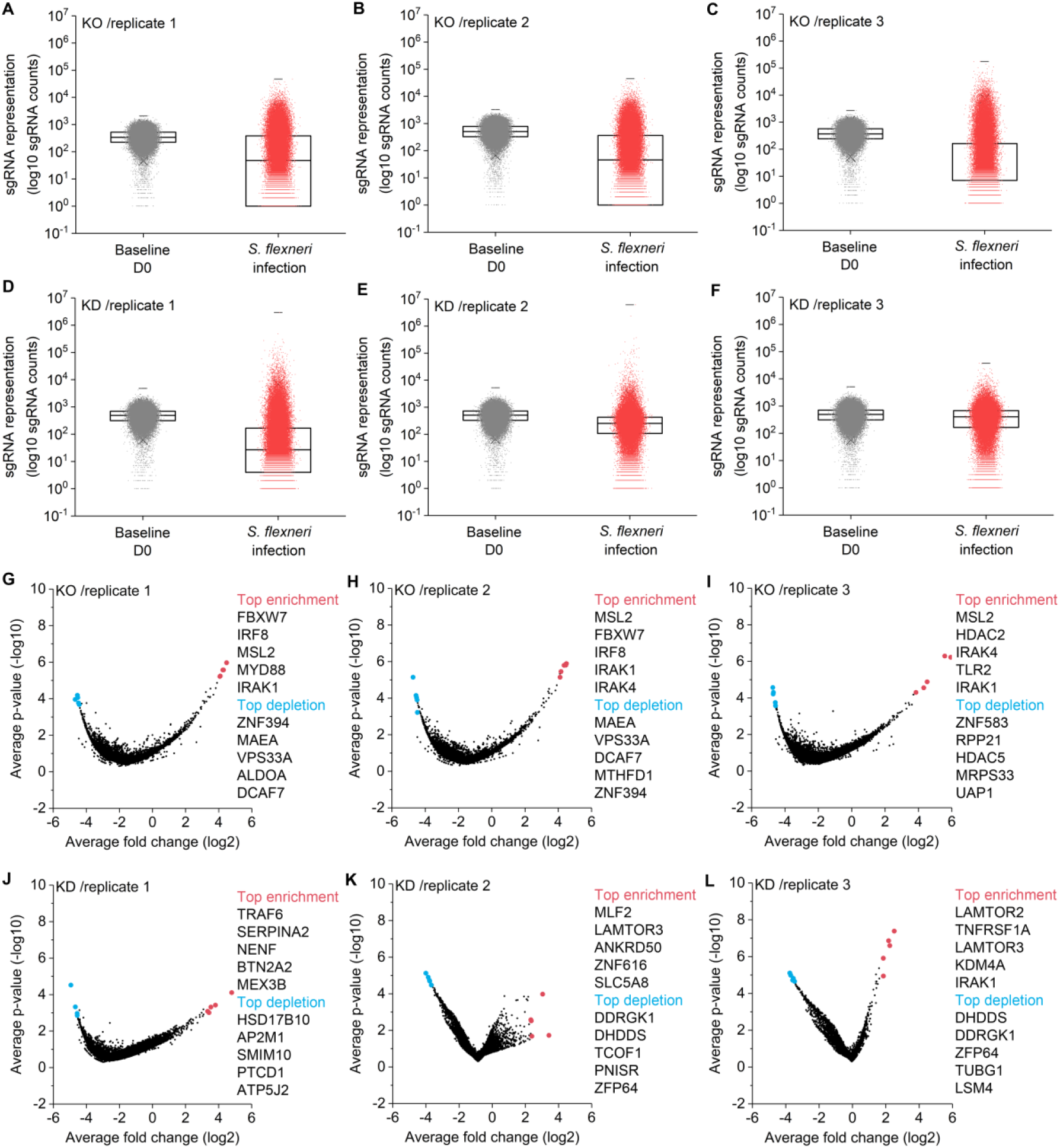
sgRNA distribution and genetic hits in genome-wide CRISPR knockout and CRISPRi screening. Related to Figure 1. (A)-(C) Distribution of individual sgRNA in control and *S. flexneri*-infected samples after CRISPR knockout screens. (D)-(F) Distribution of individual sgRNA in control and *S. flexneri*-infected samples after CRISPRi screens. Each point represents individual sgRNAs. Boxes, 25th to 75th percentile; Whiskers, 1st to 99th percentile. (G)-(I) Volcano plots from CRISPR knockout screens. (J)-(L) Volcano plots from CRISPRi screens. For each sgRNA-targeted gene, the x axis shows its enrichment (positive hits) or depletion (negative hits) post-infection, and the y axis shows statistical significance measured by p-value. Top 5 positive and negative screen hits are labeled as red and blue dots, respectively.

**Figure S3.**
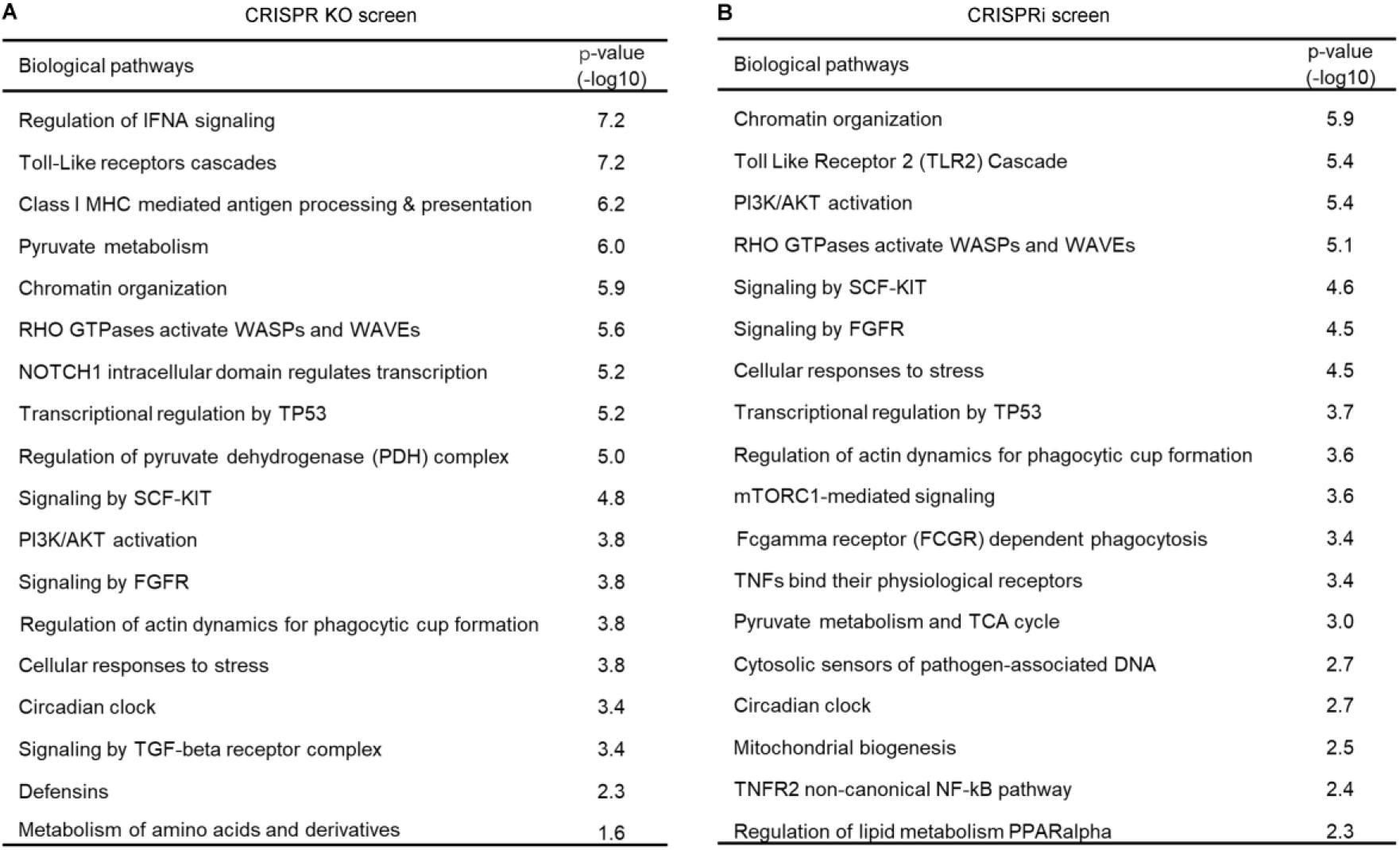
Enrichment of biological pathways identified by genome-wide CRISPR knockout and CRISPRi screens in *S. flexneri* infection of THP-1 cells. Related to Figure 2. (A) and (B) Gene enrichment analysis of positive genetic hits identified by CRISPR knockout screen (A) and CRISPRi screen (B) in *S. flexneri* infection (FDR<0.1 and log2 FC>1).

**Figure S4.**
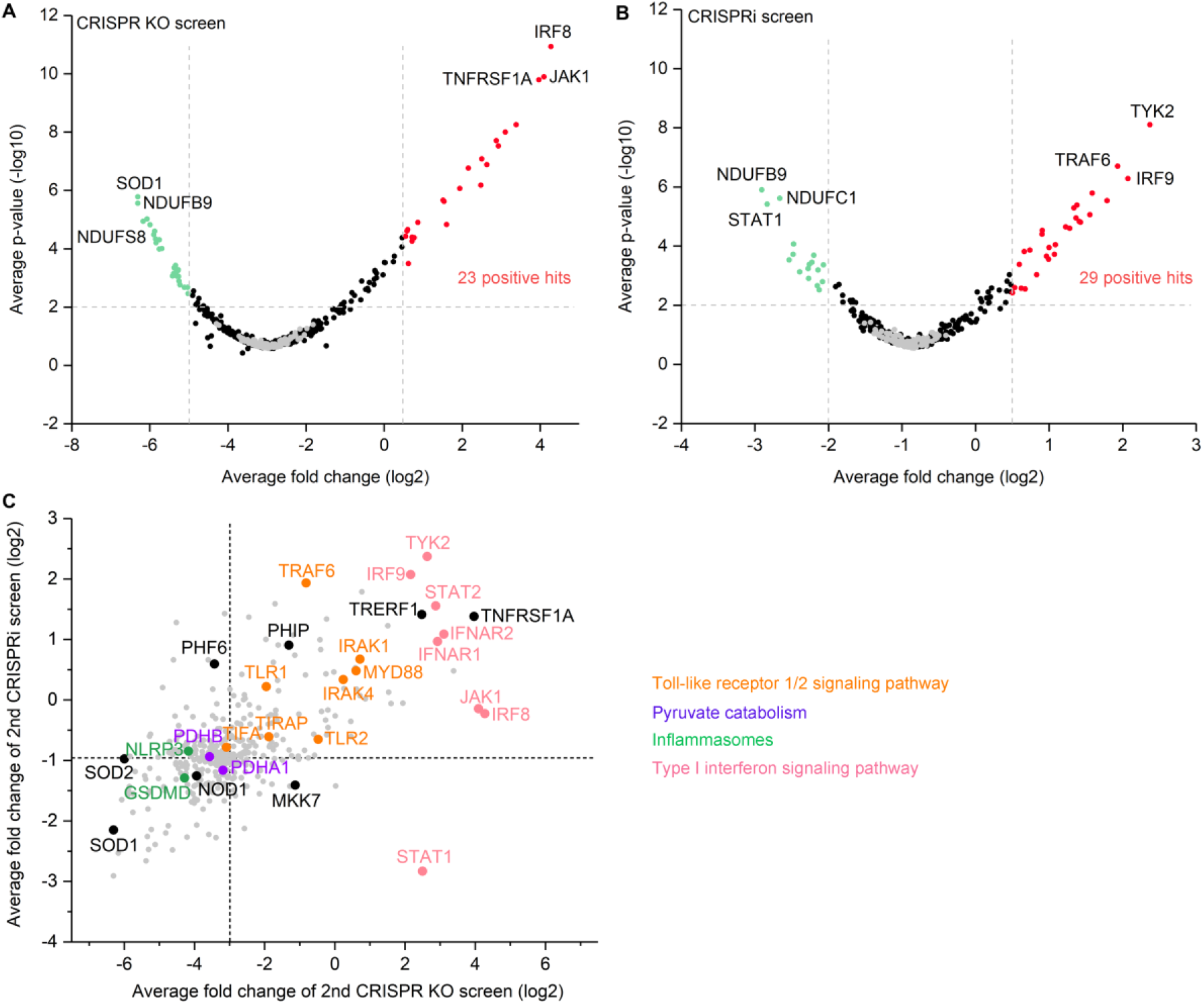
Genetic hits identified by secondary CRISPR screens in *Shigella* infection of THP-1 cells. Related to Figure 3. (A) and (B) Volcano plots from secondary CRISPR knockout (A) and CRISPRi (B) screens. For each sgRNA-targeted gene, the x axis shows its enrichment (positive hits) or depletion (negative hits) post-infection, and the y axis shows statistical significance measured by p-value. Top 3 positive and negative screen hits are labeled as red and green dots, respectively. Gray dots represent non-targeting controls. For each screen, experiments were carried out in triplicate. (C) Gene-centric visualization of average fold change for secondary screens in *S. flexneri*-infected versus non-infected host cells. Selected components of TLR 1/2 signaling pathway, pyruvate catabolism, inflammasome formation, and type I interferon signaling pathway are highlighted in orange, purple, green, and pink, respectively.

**Figure S5.**
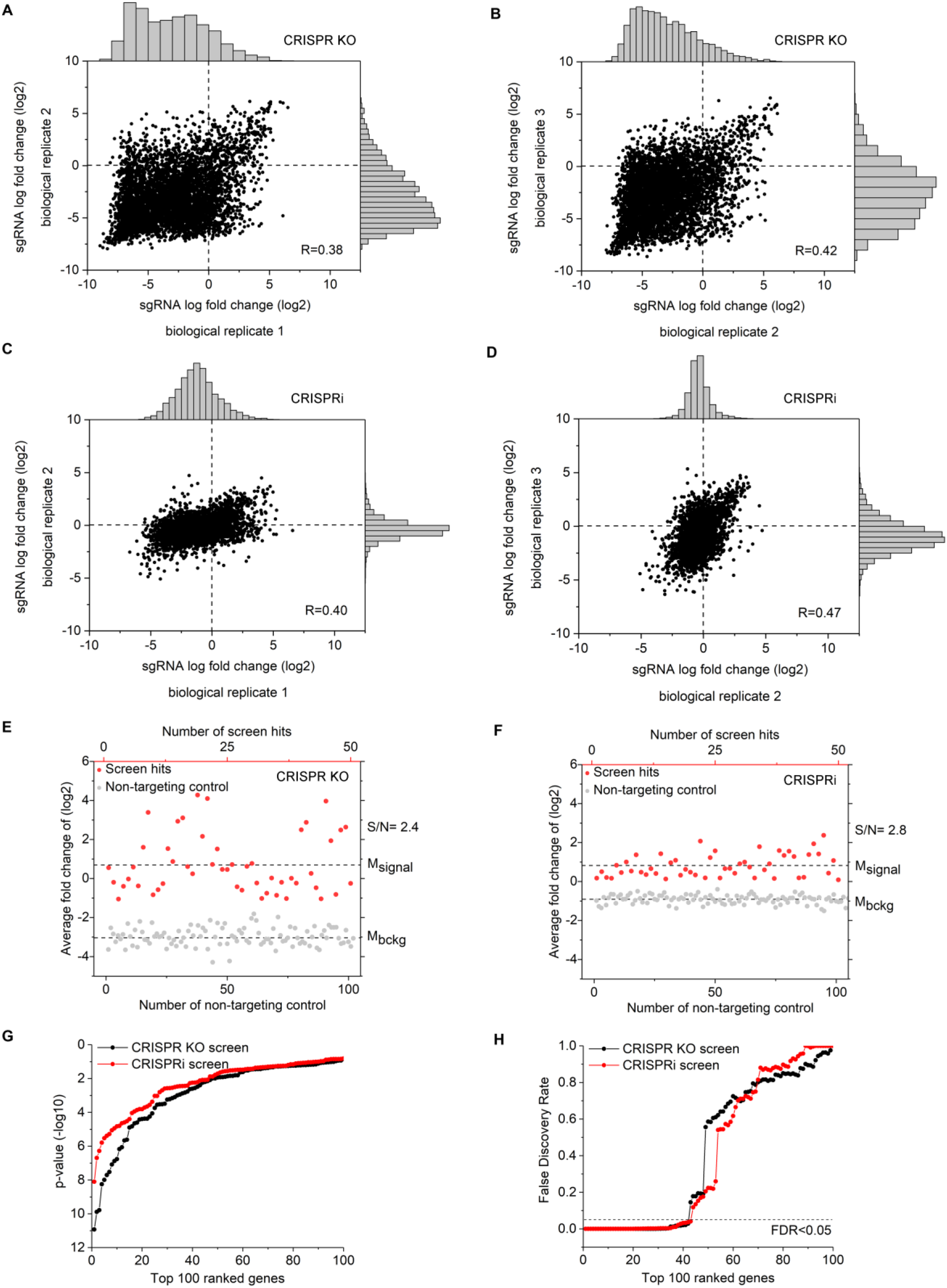
Comparison of secondary CRISPR knockout and CRISPRi screens in *S. flexneri* infection of THP-1 cells. Related to Figure 3. (A)-(D) sgRNA-level correlation of replicates in CRISPR knockout screens (A and B) and CRISPRi screens (C and D). Pearson correlation of log2 fold change values between replicates is indicated. (E) and (F) Top 50 positive genetic hits (red) and non-targeting controls (gray) identified by CRISPR knockout (E) and CRISPRi (F) in *S. flexneri* infection. Broken lines show the means of the hits (M_signal_) and non-targeting controls (M_bckg_). (G) and (H) p-value (G) and FDR (H) of top 100 positive genetic hits identified by CRISPR knockout and CRISPRi screens in *S. flexneri* infection.

**Figure S6.**
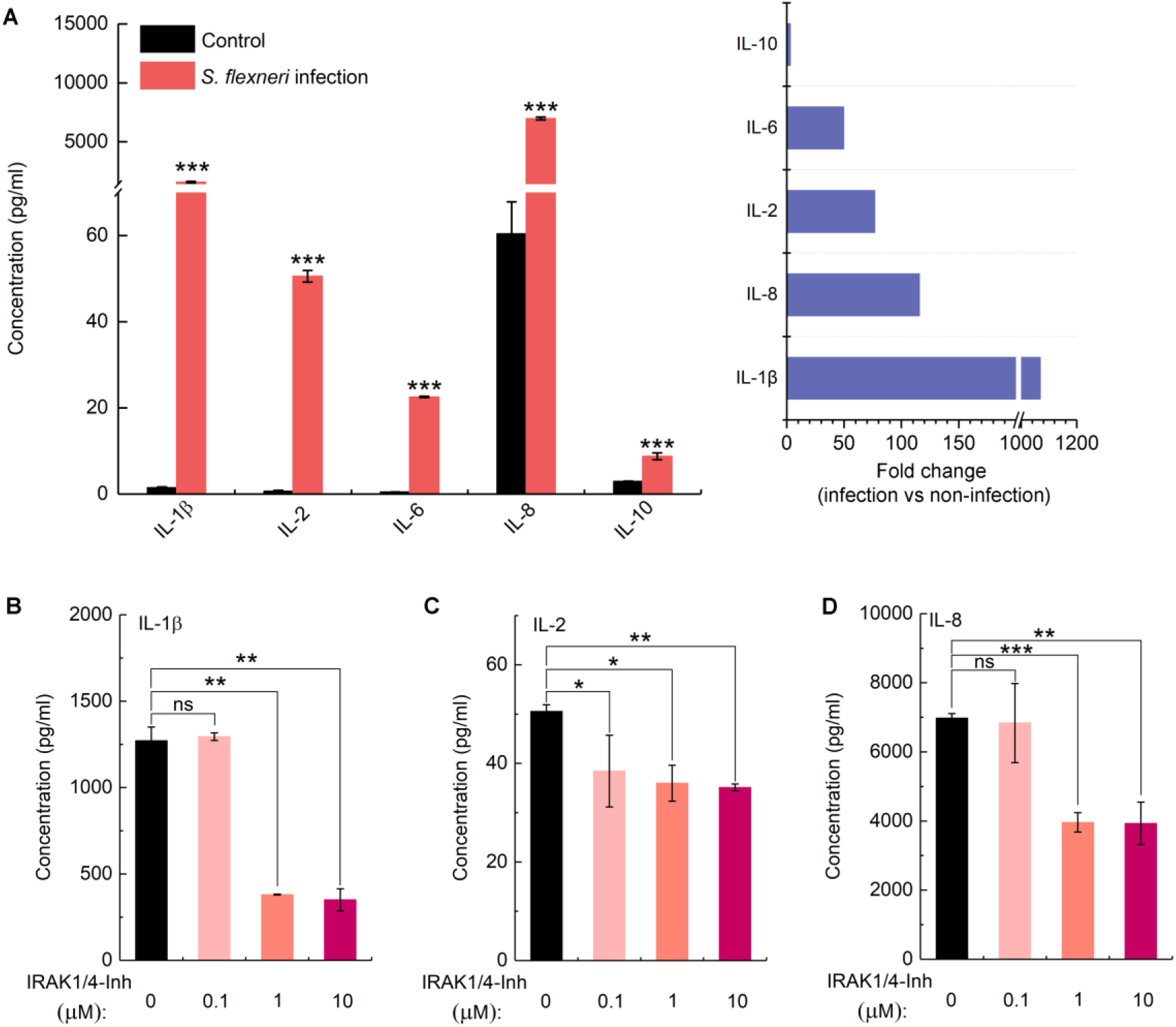
Cytokine and chemokine production in THP-1 cells in *S. flexneri* infection. Related to Figure 4. (A) Cytokine and chemokine production in THP-1 cells with or without *S. flexneri* infection. (B)-(D) Production of IL-1β (B), IL-2 (C), and IL-8 (D) in THP-1 cells in the presence of IRAKI inhibitor at different concentrations. Data represent the mean ±SD (n = 3) (twotailed unpaired Student’s *t*-test, * P<0.05 ** P<0.01 *** P<0.001; ns represents not significant).

**Figure S7.**
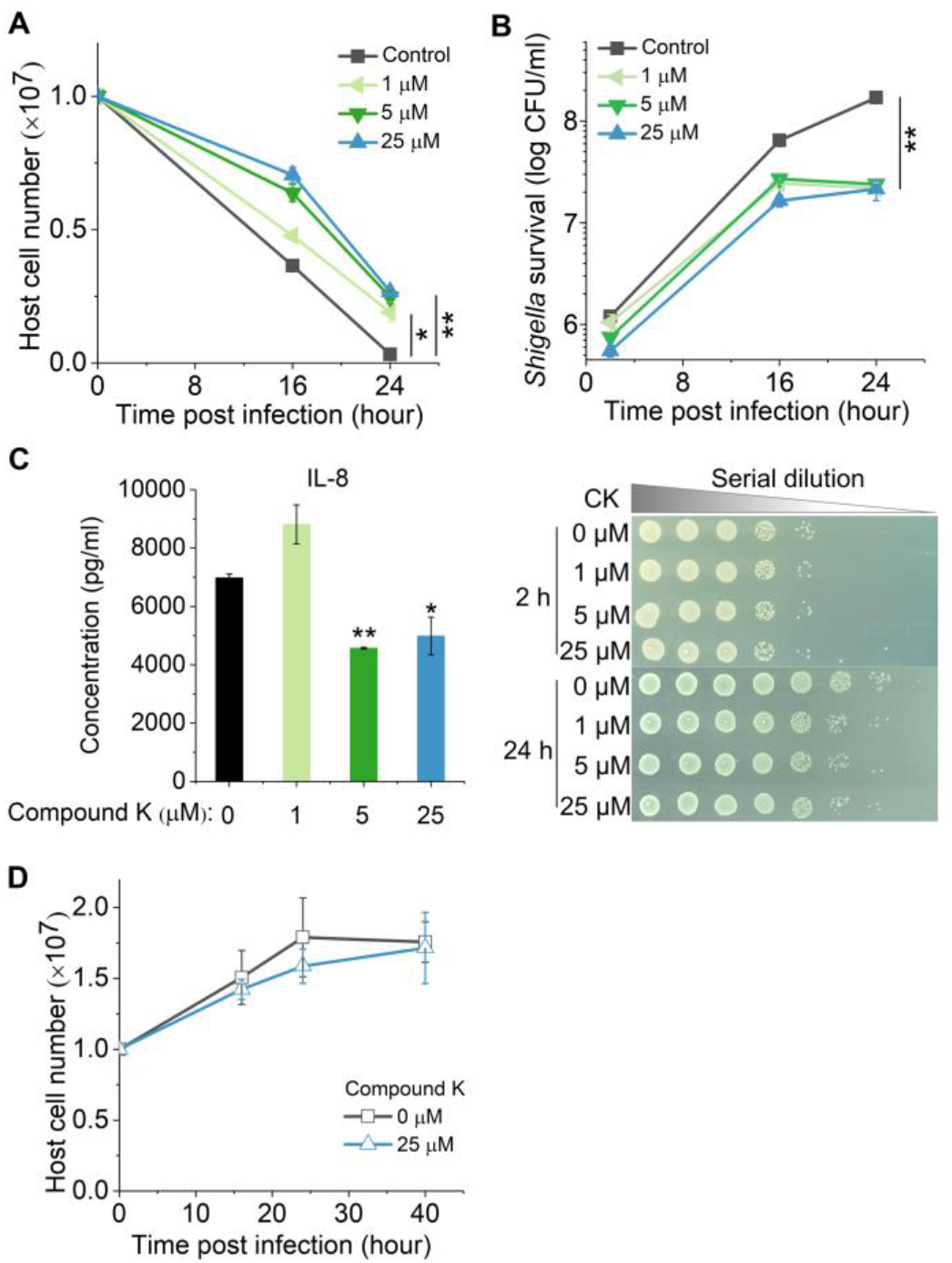
Validation of IRAK1 inhibitor Compound K in *Shigella* infection. Related to Figure 4. (A) Survival of THP-1 cells post-infection in the presence or absence of different concentrations of Compound K (CK). (B) Intracellular *S. flexneri* growth in the presence or absence of different concentrations of CK. (C) Production of infection-induced IL-8 in the presence or absence of different concentrations of CK. (D) The growth of THP-1 cells in the presence or absence of CK without *S. flexneri* infection. Data represent the mean ± SD (n =3) (two-tailed unpaired Student’s *t*-test, * P<0.05 ** P<0.01; ns represents not significant).

## STAR METHODS

### RESOURCE AVAILABILITY

#### Lead Contact

Further information and requests for resources and reagents should be directed to and will be fulfilled by the Lead Contact, Timothy K. Lu.

#### Materials Availability

Plasmids generated in this study are forthcoming to Addgene. Plasmids will be available upon request from the Lead Contact.

#### Data and Code Availability

The datasets generated during this study are available at Mendeley Data (DOI: 10.17632/xn3vv2cdnk.1). This study did not generate new code.

### EXPERIMENTAL MODEL AND SUBJECT DETAILS

#### Mammalian cell culture

The human monocyte cell line THP-1 was a gift from Jianzhu Chen (Singapore-MIT Alliance for Research and Technology). Cas9-expressing and dCas9-Krab-expressing THP-1 cell lines were constructed in previous study (Lai et al., 2020). HEK293FT cells were gifts from Asha Shekaran (Engine Biosciences). THP-1 cells were cultured in RPMI1640 (HyClone) with 10% FBS (Gibco) and Pen/Strep at 37°C with 5% CO_2_. 50 ng ml^-1^ phorbol 12-myristate 13-acetate (PMA; Sigma) was used to differentiate THP-1 cells in tissue culture treated 96-well plates (Corning). HEK293FT cells were cultured in Dulbecco’s modified Eagle’s medium (DMEM) (HyClone) supplemented with 10% FBS and Pen/Strep at 37°C with 5% CO_2_.

Frozen peripheral blood mononuclear cells were obtained by Ficoll gradient centrifugation of healthy donor leukaphereses (Research Blood Components). Primary human monocytes were isolated by CD14 positive selection (Stemcell Technologies). Monocytes were allowed to mature into macrophages on tissue culture-treated dishes using 50 ng ml^-1^ GM-CSF (BioLegend) for 6 days in RPMI1640 with 10% FBS, 10 mM HEPES, and 1× GlutaMAX (Gibco). Matured macrophages were dissociated with Accutase (Innovative Cell Technologies), counted, distributed in a 96-well plate format, and allowed to adhere overnight in the same media without GM-CSF. All incubations were performed at 37°C with 5% CO_2_.

#### Bacterial strains and growth conditions

*Shigella flexneri* M90T *ΔvirG* pCK100 (P*uhpT*::dsRed), a gift from Cecile Arrieumerlou (Institut Cochin), was grown in Lysogeny broth (LB) medium overnight at 37°C with shaking. The next day, bacteria were diluted 1:100 into 10 mL LB medium and grown to exponential phase for infection. When necessary, 100 μg ml^-1^ ampicillin was added to the growth medium.

### METHOD DETAILS

#### Reagents

IRAK1/4-Inh, oxythiamine, and ginsenoside Compound K were purchased from Sigma and used at the following concentrations: IRAK1/4-Inh, 0.1-10 μM; Oxythiamine, 0.01-1 mM; Compound K, 1-25 μM. Antibiotics in the media were at the following concentrations: 100 μg ml^-1^ ampicillin, 100 μg ml^-1^ gentamicin, 100 U ml^-1^ Penicillin-Streptomycin (Pen/Strep; Gibco).

#### *In vitro* bacterial infection

*S. flexneri* M90T *ΔvirG* pCK100 were prepared in exponential growth phase for host cell infection. THP-1 cells were infected at an MOI of 10 in complete RPMI1640 medium at indicated times. To test the combination of IRAK1 and PDHB small molecule inhibitors, THP-1 cells and primary human monocyte-derived macrophages (hMDM) were infected at an MOI of 1:10. After *Shigella* infection, THP-1 cells and hMDM were treated with 100 μg ml^-1^ gentamicin for 2 hours to kill extracellular bacteria. Subsequently, host cells were washed and maintained for the rest of the experiments. Viable THP-1 cells were counted in a hemocytometer by using trypan blue (Gibcon). Cell death of PMA-stimulated THP-1 cells and hMDM were measured by LDH assay (Takara).

#### Enumeration of intracellular bacteria in infected cells

At selected time points, 1 ml of *Shigella*-infected THP-1 cells were centrifuged and washed twice with 1×PBS and then lysed with 50 μl of 1×PBS with 1% Triton 100. PMA-stimulated THP-1 cells and hMDMs infected with *S. flexneri* were lysed with 50 μl of 1×PBS with 1% Triton 100. 10-fold serial dilutions were performed followed by plating on LB agar plates and incubated at 37°C for 24 hours. The number of viable intracellular bacteria was calculated from the counted CFU on the agar plates.

#### Pooled Genome-wide and secondary CRISPR Screens

The Brunello human CRISPR knockout pooled library was obtained from Addgene (#73178). The Dolcetto human CRISPRi pooled library was a gift from John Doench (the Broad Institute, also available on Addgene #92385). Both secondary CRISPR knockout and CRISPRi libraries, with 10 sgRNAs per gene, were designed to target the 251 genes scored in primary genome-wide screens (133 of these genes were identified as being involved in *S. flexneri* infection) and 121 genes from literature (47 of these genes were found to be involved in *S. flexneri* infection) (Lai et al., 2020). 1000 non-targeting sgRNAs were used as controls (Lai et al., 2020).

#### Lentiviral library packaging

Well dissociated HEK293FT cells were seeded at a density of 1.4×10^7^ cells per flask in a total volume of 35 ml of DMEM medium 24 h before transfection. Cells were optimal for transfection at 80-90% confluency using 7 ml of Opti-MEM, 231 μl of PLUS reagent, 210 μl of Lipofectamine 2000, and a DNA mixture of 18.2 μg of psPAX2 (Addgene #12260), 11.9 μg of pMD2.G (Addgene #12259), and 23.8 μg of library plasmid. Flasks were incubated at 37°C with 5% CO_2_ for 4 hours. The media was replaced with 35 ml DMEM medium with 10% FBS and 1% BSA. Lentivirus was harvested 2 days after the start of transfection and filtered through a 0.45 μm polyethersulfone membrane.

#### Lentivirus transduction

To ensure that only one gene was targeted in each cell, Cas9- and dCas9-Krab-expressing THP-1 cells were transduced with the pooled lentiviral CRISPR knockout and CRISPRi libraries in three biological replicates at an MOI of 0.3, respectively. To ensure that each perturbation would be fully represented and reduce spurious effects due to random genome integration in the transduced cell population, screening libraries were prepared with coverage of >500 cells per sgRNA. Lentiviral spinfection was performed by centrifuging 12-well plates at 1,000 *g* for 2 hours at 33°C with THP-1 cells grown in RPMI1640 medium with 10% FBS and 8 μg ml^-1^ of polybrene. 24 hours after lentiviral transduction, cell culture medium was replaced by RPMI1640 with 10% FBS and 2 μg ml^-1^ of puromycin for selection. Following antibiotic selection, a library coverage of >3000× was maintained for subsequent screens.

#### CRISPR screens

After puromycin selection, each CRISPR library replicate was split, one for *Shigella* infection and one used as control to verify library representation. 100 μg ml^-1^ of gentamicin was added to the cell culture to kill extracellular pathogen post-infection. Surviving host cells were harvested and pelleted by centrifugation with coverage of >500 cells per sgRNA. The pooled screens were performed as three independent replicates.

#### Genomic DNA extraction, barcode amplification, next generation sequencing (NGS), and analysis

Genomic DNA (gDNA) from live THP-1 cells was isolated using a homemade modified salt precipitation method as described previously (Chen et al., 2015). The sgRNA cassette was amplified by PCR and prepared for Illumina sequencing (HiSeq2000) as described previously (Sanson et al., 2018). The sequencing reads were deconvoluted to generate a matrix of read counts, which were then normalized under each condition by the following formula: log2 (Reads per sgRNA/total reads per condition*10^6^+1). The log2 fold change of each sgRNA was determined by comparing infected sample and uninfected samples for each biological replicate. A CRISPR screen analysis tool developed by the Genetic Perturbation Platform (GPP) at the Broad institute was used to evaluate the rank and statistical significance of genes (https://portals.broadinstitute.org/gpp/public/analysis-tools/crispr-gene-scoring). A hypergeometric distribution was used to calculate the overlap probability of screen hits between CRISPR knockout and CRISPRi screens. Tests were carried out in R package using the function phyper (q, m, n, k, lower. tail=FALSE), where q is the number of overlap genetic hits, m is the number of genetic hits identified by the CRISPR knockout screen, n is the total number of genes in the library, and k is the number of genetic hits identified by the CRISPRi screen. The GSEA tool (Mootha et al., 2003; Subramanian et al., 2005) was used to perform gene-set enrichment analysis based on genetic hits identified by CRISPR screens. Enrichment Map was used for interpretation of the biological processes (Merico et al., 2010).

#### Validation of individual sgRNAs

For each sgRNA cloning, spacer-encoding sense and antisense oligonucleotides with BsmBI-compatible overhangs were synthesized, annealed, cloned into the lentiGuide-Puro vector (Addgene #52963), and verified by sequencing (**Table S8**). Lentivirus was generated in HEK293FT cells using PLUS reagents and Lipofectamine 2000, following manufacturer’s instructions. Lentiviral transduction was performed in dCas9-Krab-expressing THP-1 cells to generate individual gene knockdown THP-1 cells. After 11 days of puromycin selection, each individual gene knockdown THP-1 cell was infected with *S. flexneri* to validate its phenotype, such as host cell survival and intracellular pathogen growth, as a top positive genetic hit identified by the CRISPR screens.

#### Cytokine quantification

Supernatants of cell cultures were collected at indicated times post-bacterial infection. Cytokine and chemokine levels in *S. flexneri*-infected supernatants were determined using Bio-plex pro human cytokine 17-plex and IFN-a2 kit (Bio-Rad) according to the manufacturer’s instructions. The results were measured by a Bio-Plex 200 system (Bio-Rad).

#### Metabolite profiling

Metabolite extraction and targeted metabolomics analyses followed the published reports with modifications (Zhong et al., 2017). Briefly, cell cultures were harvested at given time points and rapidly quenched, and metabolites were extracted using acetonitrile:methanol:water (2:2:1). After centrifugation, the supernatant was collected and evaporated to dryness in a vacuum evaporator, and the dry extracts were redissolved in 100 μL of 98:2 water/methanol for liquid chromatography-mass spectrometry (LC-MS) analysis.

The targeted LC-MS/MS analysis was performed with Agilent 1290 ultrahigh pressure liquid chromatography system coupled to a 6490 Triple Quadrupole mass spectrometer equipped with a dual-spray electrospray ionization source with Jet StreamTM (Agilent Technologies, Santa Clara, CA). Chromatographic separation of metabolites in central carbon metabolism was achieved by using Phenomenex (Torrance, CA) RezexTM ROA-Organic Acid H+ (8%) column (2.1× 100 mm, 3 μm) and the compounds were eluted at 40°C with an isocratic flow rate of 0.3 mL min^-1^ of 0.1% formic acid in water. Compounds were quantified in multiple reaction monitoring (MRM) mode. Electrospray ionization was performed in both positive and negative ion modes with the following source parameters: drying gas temperature 300°C with a flow of 10 L min^-1^, nebulizer gas pressure 40 psi, sheath gas temperature 350°C with a flow of 11 L min^-1^, nozzle voltage 500 V, and capillary voltage 4,000 V and 3,000 V for positive and negative mode, respectively. Data acquisition and processing were performed using MassHunter software (Agilent Technologies, US), and cell counts were normalized to correct variations in sample preparation.

#### Imaging

To visualize intracellular RFP-reporter *S. flexneri* M90T *ΔvirG* (P*uhpT*::dsRed), infected THP-1 cells were directly observed under a confocal fluorescence microscope (Zeiss LSM 700).

### QUANTIFICATION AND STATISTICAL ANALYSIS

Excel were used for all statistical analysis. All of the statistical details of experiments can be found in the corresponding figure legends.

## SUPPLEMENTAL INFORMATION

**Table S1.** Positive genetic hits associated biological pathways from the genome-wide CRISPR-Cas9 knockout screen in *S. flexneri* infection. Related to Figure 2.

**Table S2.** Positive genetic hits associated biological pathways from the genome-wide CRISPRi screen in *S. flexneri* infection. Related to Figure 2.

**Table S3.** Positive genetic hits from the genome-wide CRISPR-Cas9 knockout screen in *S. flexneri* infection. Related to Figure 2.

**Table S4.** Positive genetic hits from the genome-wide CRISPRi screen in *S. flexneri* infection. Related to Figure 2.

**Table S5.** Positive genetic hits from secondary CRISPR-Cas9 knockout screen in *S. flexneri* infection. Related to Figure 3.

**Table S6**. Positive genetic hits from secondary CRISPRi screen in *S. flexneri* infection. Related to Figure 3.

**Table S7**. Average log2-fold change of host cell targets of *S. flexneri* virulence factors in genome-wide CRISPR screens. Related to Figure 2.

**Table S8**. Positive genetic hits validated in *S. flexneri* infection. Related to Figure 4 and STAR Methods.

## References

Ashida, H., Kim, M., and Sasakawa, C. (2014). Manipulation of the host cell death pathway by *Shigella*. Cell Microbiol 16, 1757–1766.

Ashida, H., Mimuro, H., and Sasakawa, C. (2015). *Shigella* manipulates host immune responses by delivering effector proteins with specific roles. Front Immunol 6, 219.

Ashida, H., Ogawa, M., Kim, M., Suzuki, S., Sanada, T., Punginelli, C., Mimuro, H., and Sasakawa, C. (2011). *Shigella* deploy multiple countermeasures against host innate immune responses. Curr Opin Microbiol 14, 16–23.

Bergounioux, J., Elisee, R., Prunier, A.L., Donnadieu, F., Sperandio, B., Sansonetti, P., and Arbibe, L. (2012). Calpain activation by the *Shigella flexneri* effector VirA regulates key steps in the formation and life of the bacterium’s epithelial niche. Cell Host Microbe 11, 240–252.

Carneiro, L.A., Travassos, L.H., Soares, F., Tattoli, I., Magalhaes, J.G., Bozza, M.T., Plotkowski, M.C., Sansonetti, P.J., Molkentin, J.D., Philpott, D.J., et al. (2009). *Shigella* induces mitochondrial dysfunction and cell death in nonmyleoid cells. Cell Host Microbe 5, 123–136.

Chen, S., Sanjana, N.E., Zheng, K., Shalem, O., Lee, K., Shi, X., Scott, D.A., Song, J., Pan, J.Q., Weissleder, R., et al. (2015). Genome-wide CRISPR screen in a mouse model of tumor growth and metastasis. Cell 160, 1246–1260.

Daniloski, Z., Jordan, T.X., Wessels, H.-H., Hoagland, D.A., Kasela, S., Legut, M., Maniatis, S., Mimitou, E.P., Lu, L., Geller, E., et al. (2020). Identification of required host factors for SARS-CoV-2 infection in human cells. Cell. in press

Doench, J.G., Fusi, N., Sullender, M., Hegde, M., Vaimberg, E.W., Donovan, K.F., Smith, I., Tothova, Z., Wilen, C., Orchard, R., et al. (2016). Optimized sgRNA design to maximize activity and minimize off-target effects of CRISPR-Cas9. Nat Biotechnol 34, 184–191.

Ferrari, M.L., Malarde, V., Grassart, A., Salavessa, L., Nigro, G., Descorps-Declere, S., Rohde, J.R., Schnupf, P., Masson, V., Arras, G., et al. (2019). *Shigella* promotes major alteration of gut epithelial physiology and tissue invasion by shutting off host intracellular transport. Proc Natl Acad Sci U S A 116, 13582–13591.

Garcia-Weber, D., Dangeard, A.S., Cornil, J., Thai, L., Rytter, H., Zamyatina, A., Mulard, L.A., and Arrieumerlou, C. (2018). ADP-heptose is a newly identified pathogen-associated molecular pattern of *Shigella flexneri*. EMBO Rep 19. e46943

Gaudet, R.G., Guo, C.X., Molinaro, R., Kottwitz, H., Rohde, J.R., Dangeard, A.S., Arrieumerlou, C., Girardin, S.E., and Gray-Owen, S.D. (2017). Innate recognition of intracellular bacterial growth is driven by the TIFA-dependent cytosolic surveillance pathway. Cell Rep 19, 1418–1430.

Gu, B., Cao, Y., Pan, S., Zhuang, L., Yu, R., Peng, Z., Qian, H., Wei, Y., Zhao, L., Liu, G., et al. (2012). Comparison of the prevalence and changing resistance to nalidixic acid and ciprofloxacin of *Shigella* between Europe-America and Asia-Africa from 1998 to 2009. Int J Antimicrob Agents 40, 9–17.

He, S., Liang, Y., Shao, F., and Wang, X. (2011). Toll-like receptors activate programmed necrosis in macrophages through a receptor-interacting kinase-3-mediated pathway. Proc Natl Acad Sci U S A 108, 20054–20059.

Kentner, D., Martano, G., Callon, M., Chiquet, P., Brodmann, M., Burton, O., Wahlander, A., Nanni, P., Delmotte, N., Grossmann, J., et al. (2014). *Shigella* reroutes host cell central metabolism to obtain high-flux nutrient supply for vigorous intracellular growth. Proc Natl Acad Sci USA 111, 9929–9934.

Khalil, I.A., Troeger, C., Blacker, B.F., Rao, P.C., Brown, A., Atherly, D.E., Brewer, T.G., Engmann, C.M., Houpt, E.R., Kang, G., et al. (2018). Morbidity and mortality due to *Shigella* and enterotoxigenic *Escherichia coli* diarrhoea: the global burden of disease study 1990-2016. Lancet Infect Dis 18, 1229–1240.

Killackey, S.A., Sorbara, M.T., and Girardin, S.E. (2016). Cellular aspects of *Shigella* pathogenesis: focus on the manipulation of host cell processes. Front Cell Infect Microbiol 6, 38.

Kobayashi, T., Ogawa, M., Sanada, T., Mimuro, H., Kim, M., Ashida, H., Akakura, R., Yoshida, M., Kawalec, M., Reichhart, J.M., et al. (2013). The *Shigella* OspC3 effector inhibits caspase-4, antagonizes inflammatory cell death, and promotes epithelial infection. Cell Host Microbe 13, 570–583.

Kotloff, K.L., Riddle, M.S., Platts-Mills, J.A., Pavlinac, P., and Zaidi, A.K.M. (2018). Shigellosis. Lancet 391, 801–812.

Lai, Y., Babunovic, G.H., Cui, L., Dedon, P.C., Doench, J.G., Fortune, S.M., and Lu, T.K. (2020). Illuminating host-mycobacterial interactions with genome-wide CRISPR knockout and CRISPRi screens. Cell Syst 11, 239–251 e237.

Liu, X., Zhang, Z., Ruan, J., Pan, Y., Magupalli, V.G., Wu, H., and Lieberman, J. (2016). Inflammasome-activated gasdermin D causes pyroptosis by forming membrane pores. Nature 535, 153–158.

Merico, D., Isserlin, R., Stueker, O., Emili, A., and Bader, G.D. (2010). Enrichment map: a network-based method for gene-set enrichment visualization and interpretation. PLoS One 5, e13984.

Miao, E.A., Leaf, I.A., Treuting, P.M., Mao, D.P., Dors, M., Sarkar, A., Warren, S.E., Wewers, M.D., and Aderem, A. (2010). Caspase-1-induced pyroptosis is an innate immune effector mechanism against intracellular bacteria. Nat Immunol 11, 1136–1194.

Milivojevic, M., Dangeard, A.S., Kasper, C.A., Tschon, T., Emmenlauer, M., Pique, C., Schnupf, P., Guignot, J., and Arrieumerlou, C. (2017). ALPK1 controls TIPA/TRAP6-dependent innate immunity against heptose-1,7-bisphosphate of gram-negative bacteria. PLoS Pathog 13, e1006224.

Mootha, V.K., Lindgren, C.M., Eriksson, K.F., Subramanian, A., Sihag, S., Lehar, J., Puigserver, P., Carlsson, E., Ridderstrale, M., Laurila, E., et al. (2003). PGC-1alpha-responsive genes involved in oxidative phosphorylation are coordinately downregulated in human diabetes. Nat Genet 34, 267–273.

Niebuhr, K., Giuriato, S., Pedron, T., Philpott, D.J., Gaits, F., Sable, J., Sheetz, M.P., Parsot, C., Sansonetti, P.J., and Payrastre, B. (2002). Conversion of PtdIns(4,5)P-2 into PtdIns(5)P by the *S. flexneri* effector IpgD reorganizes host cell morphology. EMBO J 21, 5069–5078.

Oliveira-Nascimento, L., Massari, P., and Wetzler, L.M. (2012). The role of TLR2 in infection and immunity. Front Immunol 3, 79.

Pendaries, C., Tronchere, H., Arbibe, L., Mounier, J., Gozani, O., Cantley, L., Fry, M.J., Gaits-Iacovoni, F., Sansonetti, P.J., and Payrastre, B. (2006). PtdIns5P activates the host cell PI3-kinase/Akt pathway during *Shigella flexneri* infection. EMBO J 25, 1024–1034.

Ramel, D., Lagarrigue, F., Pons, V., Mounier, J., Dupuis-Coronas, S., Chicanne, G., Sansonetti, P.J., Gaits-Iacovoni, F., Tronchere, H., and Payrastre, B. (2011). *Shigella flexneri* infection generates the lipid PI5P to alter endocytosis and prevent termination of EGFR signaling. Sci Signal 4, ra61.

Runyen-Janecky, L.J., and Payne, S.M. (2002). Identification of chromosomal *Shigella flexneri* genes induced by the eukaryotic intracellular environment. Infect Immun 70, 4379–4388.

Sanson, K.R., Hanna, R.E., Hegde, M., Donovan, K.F., Strand, C., Sullender, M.E., Vaimberg, E.W., Goodale, A., Root, D.E., Piccioni, F., et al. (2018). Optimized libraries for CRISPR-Cas9 genetic screens with multiple modalities. Nat Commun 9, 5416.

Senerovic, L., Tsunoda, S.P., Goosmann, C., Brinkmann, V., Zychlinsky, A., Meissner, F., and Kolbe, M. (2012). Spontaneous formation of IpaB ion channels in host cell membranes reveals how *Shigella* induces pyroptosis in macrophages. Cell Death Dis 3, e384.

Spiller, S., Elson, G., Ferstl, R., Dreher, S., Mueller, T., Freudenberg, M., Daubeuf, B., Wagner, H., and Kirschning, C.J. (2008). TLR4-induced IFN-gamma production increases TLR2 sensitivity and drives Gram-negative sepsis in mice. J Exp Med 205, 1747–1754.

Subramanian, A., Tamayo, P., Mootha, V.K., Mukherjee, S., Ebert, B.L., Gillette, M.A., Paulovich, A., Pomeroy, S.L., Golub, T.R., Lander, E.S., et al. (2005). Gene set enrichment analysis: a knowledge-based approach for interpreting genome-wide expression profiles. Proc Natl Acad Sci U S A 102, 15545–15550.

Suzuki, S., Franchi, L., He, Y., Munoz-Planillo, R., Mimuro, H., Suzuki, T., Sasakawa, C., and Nunez, G. (2014a). *Shigella* type III secretion protein MxiI is recognized by Naip2 to induce Nlrc4 inflammasome activation independently of Pkcdelta. PLoS Pathog 10, e1003926.

Suzuki, S., Mimuro, H., Kim, M., Ogawa, M., Ashida, H., Toyotome, T., Franchi, L., Suzuki, M., Sanada, T., Suzuki, T., et al. (2014b). *Shigella* IpaH7.8 E3 ubiquitin ligase targets glomulin and activates inflammasomes to demolish macrophages. Proc Natl Acad Sci U S A 111, E4254–E4263.

Suzuki, T., Franchi, L., Toma, C., Ashida, H., Ogawa, M., Yoshikawa, Y., Mimuro, H., Inohara, N., Sasakawa, C., and Nunez, G. (2007). Differential regulation of caspase-1 activation, pyroptosis, and autophagy via Ipaf and ASC in *Shigella*-infected macrophages. Plos Pathog 3, 1082–1091.

Waligora, E.A., Fisher, C.R., Hanovice, N.J., Rodou, A., Wyckoff, E.E., and Payne, S.M. (2014). Role of intracellular carbon metabolism pathways in *Shigella flexneri* virulence. Infect Immun 82, 2746–2755.

Wassef, J.S., Keren, D.F., and Mailloux, J.L. (1989). Role of M-cells in initial antigen uptake and in ulcer formation in the rabbit intestinal loop model of Shigellosis. Infect Immun 57, 858–863.

Wee, Z.N., Yatim, S.M.J., Kohlbauer, V.K., Feng, M., Goh, J.Y., Bao, Y., Lee, P.L., Zhang, S., Wang, P.P., and Lim, E. (2015). IRAK1 is a therapeutic target that drives breast cancer metastasis and resistance to paclitaxel. Nat Commun 6, 1–16.

Wei, J., Alfajaro, M.M., DeWeirdt, P.C., Hanna, R.E., Lu-Culligan, W.J., Cai, W.L., Strine, M.S., Zhang, S.M., Graziano, V.R., Schmitz, C.O., et al. (2020). Genome-wide CRISPR screens reveal host factors critical for SARS-CoV-2 infection. Cell. in press

Wiersinga, W.J., Wieland, C.W., Dessing, M.C., Chantratita, N., Cheng, A.C., Limmathurotsakul, D., Chierakul, W., Leendertse, M., Florquin, S., de Vos, A.F., et al. (2007). Toll-like receptor 2 impairs host defense in gram-negative sepsis caused by *Burkholderia pseudomallei* (Melioidosis). PLoS Med 4, e248.

Willingham, S.B., Bergstralh, D.T., O’Connor, W., Morrison, A.C., Taxman, D.J., Duncan, J.A., Barnoy, S., Venkatesan, M.I., Flavell, R.A., Deshmukh, M., et al. (2007). Microbial pathogen-induced necrotic cell death mediated by the inflammasome components CIAS1/Cryopyrin/NLRP3 and ASC. Cell Host Microbe 2, 147–159.

World Health Organization and UNICEF (2013). Ending preventable child deaths from pneumonia and diarrhoea by 2025: The integrated global action plan for pneumonia and diarrhoea (GAPPD).

Zhong, W., Cui, L., Goh, B.C., Cai, Q., Ho, P., Chionh, Y.H., Yuan, M., Sahili, A.E., Fothergill-Gilmore, L.A., Walkinshaw, M.D., et al. (2017). Allosteric pyruvate kinase-based “logic gate” synergistically senses energy and sugar levels in *Mycobacterium tuberculosis*. Nat Commun 8, 1986.

Zhou, P., She, Y., Dong, N., Li, P., He, H.B., Borio, A., Wu, Q.C., Lu, S., Ding, X.J., Cao, Y., et al. (2018). Alpha-kinase 1 is a cytosolic innate immune receptor for bacterial ADP-heptose. Nature 561, 122–126.

