## Supplementary material for "High-throughput CRISPR screens to dissect macrophage-*Shigella* interactions": Table S7

**Table S7. Average log2-fold change of host cell targets of *S. flexneri* virulence factors in genome-wide CRISPR screens. Related to Figure 2.**

|  |  |  | **Genome-wide CRISPR knockout screen** | | **Genome-wide CRISPRi screen** | |
| --- | --- | --- | --- | --- | --- | --- |
| ***S. flexneri* effector** | **Host cell target(s)** | **Gene Symbol** | **Average LFC** | **Individual LFCs** | **Average LFC** | **Individual LFCs** |
| IpaA | Vinculin | VCL | -0.804813882 | -2.3;-0.51;-0.28;-0.14 | -2.032136652 | -3.36;-2.45;-0.29 |
|  | β1-integrins | ITGB1 | -0.556763829 | -1.3;-1.21;-1.04;1.31 | -2.912450723 | -3.89;-2.5;-2.34 |
| IpaB | CD44 | CD44 | -0.59988365 | -2.16;-0.8;0.26;0.3 | -1.927042458 | -5.02;-1.3;0.54 |
|  | caspase-1 | CASP1 | -2.651448972 | -2.89;-2.87;-2.82;-2.03 | -2.254048514 | -5.89;-1.84;0.97 |
| IpaC | Actin | ACTB | -3.204446224 | -3.7;-3.66;-3.02;-2.44 | -5.040930607 | -5.61;-5.55;-3.96 |
|  | β-catenin | CTNNB1 | 0.010257743 | -0.71;-0.54;-0.45;1.74 | -1.167230773 | -2.7;-1.07;0.27 |
| IpaH7.8 | glomulin | GLMN | -3.88340169 | -4.04;-3.95;-3.89;-3.65 | -4.328378055 | -5.42;-4.71;-2.86 |
| IpaH9.8 | Splicing factor U2AF | U2AF2 | -1.989909868 | -2.51;-2.03;-1.85;-1.58 | -2.354125386 | -3.48;-2.59;-1.0 |
| IcsAa (VirG) | N-WASP | WASL | -1.745884856 | -3.49;-1.69;-1.04;-0.76 | -3.4434207 | -4.17;-3.68;-2.49 |
| IpgB1 | ELMO protein | ELMO1 | -1.124295472 | -1.95;-1.35;-1.17;-0.03 | -3.310244532 | -4.21;-2.97;-2.75 |
| OspG | ubiquitin-conjugating enzymes | UBE2D1 | -3.23538102 | -3.39;-3.34;-3.27;-2.93 | -3.791234235 | -4.5;-3.89;-2.98 |
| OspF | p38 | MAPK14 | -2.322200145 | -2.94;-2.63;-2.34;-1.39 | -1.808376092 | -3.36;-2.49;0.43 |
| VirA | α-Tubulin | TUBA1A | -1.126597401 | -3.54;-1.71;-0.53;1.28 | -3.568511408 | -5.74;-4.14;-0.83 |
| MxiI | NLR Family Apoptosis Inhibitory Protein | NAIP | -1.293464953 | -3.53;-1.19;-1.12;0.67 | -3.766261951 | -4.7;-3.42;-3.18 |
