## Supplementary material for "High-throughput CRISPR screens to dissect macrophage-*Shigella* interactions": Table S8

**Table S8. Positive genetic hits validated in S. flexneri infection. Related to Figure 4 and STAR Methods.**

| **Bacterial infection** | **Pathways** | **Screen hits** | **sgRNA target sequence** |
| --- | --- | --- | --- |
| *S. flexneri* M90T infection | Toll-like receptors cascades | TRAF6 | GCGGCCAGCGAAGGTGGCGA |
|  |  | IRAK1 | GGACCTGCCGGGGCCTCTCA |
|  |  | MYD88 | CCTCCTGCAGCCATGGCGGG |
|  | Pyruvate metabolism | PDHB | GGGAGGTAGGCAGCAGCGCG |
|  |  | DLAT | GGGGGGTTGGTGGCACTATG |
|  |  | CS | GCCGCGCCGACGGGTTGACA |
|  | Antimicrobial peptide production | TRERF1 | GGAGAGTTTGGAGTTGCTTG |
|  | Type I IFN signaling pathway | TYK2 | TCAAGCGCAGCCAGTCCCCG |
|  | Transcriptional regulation of TP53 | TP73 | AAGGGGACGCAGCGAAACCG |
|  | Apoptosis Modulation and Signaling | TNFRSF1A | CTGGACTGAGGCTCCAGTTC |
|  | PIP3 activates AKT signaling | MKK7 | TTGTCTGCCGGACTGACGGG |
|  | Unknown | PHIP | TGAATGGTGGAGCCGAAGCT |
|  |  | PHF6 | TCCAGCAGTGCCTGAGAGCG |
| Non-targeting control |  | NC80 | ACATGTGGCTCCGCCCACAG |
|  |  | NC135 | AGATCTGCTCCATGTCACCA |
